# Lhcf2 in the peripheral antenna is essential for non-photochemical quenching and Lhcx1 accumulation in the diatom *Chaetoceros gracilis*

**DOI:** 10.64898/2026.01.17.700047

**Authors:** Jian Xing, Minoru Kumazawa, Kentaro Ifuku

**Author notes:** Correspondence: Kentaro Ifuku.

## Abstract

Photosynthetic organisms must continuously balance efficient light harvesting with protection against excess excitation energy, a challenge met by nonphotochemical quenching (NPQ). Although the molecular components involved in NPQ have been extensively studied, how the essential energy-quenching site is assembled remains poorly understood, particularly in marine diatoms. Here, we show that in the centric diatom *Chaetoceros gracilis*, which belongs to one of the most abundant and diverse genera in marine phytoplankton, the light-harvesting complex protein Lhcf2 is required for energy-dependent NPQ (qE). Targeted knockout of Lhcf2 abolished qE by preventing the stable accumulation of Lhcx1, the canonical NPQ effector in this species. Lhcf2 localizes to the peripheral antenna system and associates with Lhcx1 in a higher-order complex suggested by biochemical and functional analyses. In contrast, other established NPQ-related factors, including the trans-thylakoid proton gradient and the accumulation of diatoxanthin, were not affected by the loss of Lhcf2. These results identify a non-Lhcx-type light-harvesting complex protein as an essential structural component for qE-NPQ and establish a general design principle for the cooperative assembly of photoprotective energy-quenching sites in eukaryotic photosynthesis, with implications for marine carbon fixation.

**Significance Statement:** Photosynthetic organisms must balance efficient light harvesting with protection against excess excitation energy. Nonphotochemical quenching (NPQ) is a conserved photoprotective mechanism, yet how the essential energy-quenching site is assembled remains unclear. Here, we show that in the marine diatom *Chaetoceros gracilis*, the peripheral light-harvesting complex protein Lhcf2 is strictly required for energy-dependent NPQ by enabling the stable accumulation and functional organization of the canonical NPQ effector Lhcx1. Our results reveal that photoprotective energy dissipation depends on cooperative interactions between distinct classes of light-harvesting complex proteins rather than on a single specialized factor. This study establishes a general design principle for NPQ machinery in eukaryotic photosynthesis, with implications for marine carbon fixation under fluctuating light environments.

## Introduction

Photosynthetic organisms use light to generate a photo-current, which drives the assimilation of CO_2_ into organic matter, thereby feeding the entire biosphere and fundamentally altering the biogeography of our planet. Around 1.5 billion years ago, an ancestor of extant cyanobacteria was engulfed by a heterotrophic unicellular eukaryote, causing endosymbiosis, which further resulted in the production of a novel and vertically inherited eukaryotic organelle, the plastid (Falkowski et al. 2004; Katz et al. 2004; Parfrey et al. 2011; Keeling 2013). Land plants, green algae, and red algae derive their plastids directly from this event. In contrast, diatoms, which form a major proportion in the contemporary oceans and account for approximately half of global carbon fixation, harbor a complex “red plastid” acquired through secondary endosymbiosis of eukaryotic red algae (Pierella Karlusich et al. 2020, 2024).

All photosynthetic organisms, including diatoms with chimeric and divergent genetic backgrounds, have evolved to adapt to ever-changing light conditions. The variability in light environments encompasses spectral and intensity dimensions. The abundance of blue and green wavelengths in the ocean leads to distinct pigment compositions and coloration in marine diatoms compared to those of terrestrial organisms. Besides, light intensity is dynamic and functions as an essential energy source (Demmig-Adams and Adams 2006; Bassi and Dall’Osto 2021). In low-light (LL) conditions, optimal photosynthesis is maintained by balancing the photons absorbed by light-harvesting complex (LHC) with downstream carbon fixation. However, in high light (HL) conditions, in which the photons absorbed exceed the requirements of downstream carbon fixation processes, the accumulation of excess electrons and chlorophylls in an overexcited state can lead to the formation of reactive oxygen species (ROS) which induce cellular damage (Niyogi 1999; Takahashi and Badger 2011). To prevent damage induced by ROS, photosynthetic organisms across all phylogenetic groups have evolved a range of protective strategies.

Nonphotochemical quenching (NPQ), a conserved mechanism converts surplus excitation energy into heat (Müller et al. 2001; Ruban 2016; Bassi and Dall’Osto 2021; Van Amerongen and Croce 2025). Energy-dependent NPQ (qE) relies on the actions of trans-thylakoid proton gradient (ΔpH), de-epoxidized xanthophyll carotenoids, and effector proteins. The initiation of qE is driven by the establishment of ΔpH across the thylakoid membrane upon excess light exposure. The increased ΔpH activates xanthophyll-cycle enzymes. In diatoms, this leads to the rapid conversion of diadinoxanthin (Ddx) into diatoxanthin (Dtx) by de-epoxidase activation (Olaizola et al. 1994; Lavaud et al. 2002; Goss et al. 2006). Concurrently, the increased ΔpH leads to acidification of the thylakoid lumen, detected by specific residues on effector proteins. These proteins are PsbS and LHC stress-related (Lhcsr) protein in plants and green algae, respectively (Peers et al. 2009). PsbS is a four-helix LHC-superfamily protein that does not bind pigments or not as tightly as other LHC family proteins. As it is capable of sensing the lumen pH, PsbS induces NPQ by promoting conformational changes in nearby LHCs. The exact molecular mechanism remains unclear (Li et al. 2000; Dominici et al. 2002; Bonente et al. 2008; Wilk et al. 2013; Fan et al. 2015; Correa-Galvis et al. 2016). Lhcsr is structurally distinct from the PsbS in plant, predicted to contain three transmembrane α-helices and lumen-exposed amphipathic helices (Peers et al. 2009; Bonente et al. 2011). Lhcsr3 binds pigments and switches from a light-harvesting state to a quenching state upon protonation at its lumenal side. This dual capacity establishes Lhcsr3 as a ΔpH sensor and a direct quencher (Liguori et al. 2017; Tian et al. 2019). Despite the distant evolutionary relationship between green algae and diatoms, Lhcsr protein homologs have been identified in diatoms, where they are referred to as Lhcx proteins (Bailleul et al. 2010).

In diatoms, the importance of the Lhcsr-related Lhcx1 proteins for NPQ has been confirmed by knocking out or silencing of the *Lhcx1* genes in *Phaeodactylum tricornutum* (*P. tricornutum*), *Cyclotella meneghiniana* (*C. meneghiniana*) and *C. gracilis* (Bailleul et al. 2010; Ghazaryan et al. 2016; Buck et al. 2019; Kumazawa et al. 2025). Unlike green algae, in which Lhcsr expression is strongly HL-dependent and remains minimal under LL conditions, diatoms constitutively express a certain amount of the Lhcx1 protein (Tian et al. 2019; Giovagnetti et al. 2022). This unique constitutive expression allows diatoms to swiftly activate NPQ in response to abrupt changes in light even when grown under LL conditions (Zhang et al. 2024). NPQ, initially discovered in plants, has been extensively researched in terrestrial plants and green algae, yielding substantial progress in the elucidation of its mechanisms (Demmig-Adams et al. 2014). Diatoms lack the state transition mechanism and maintain the basal expression of Lhcx1 proteins even under LL conditions, enabling rapid activation of qE (Goss and Lepetit. 2015). Thus, most of the NPQ observed in LL-grown diatoms, without state transition-related NPQ (qT) and with minimal photoinhibition-related NPQ (qI), is largely restricted to qE. This simplifies their photoprotective response, rendering it naturally amenable to a reductionist approach for dissecting constraints and regulatory networks governing NPQ (Croteau et al. 2025). Nevertheless, the NPQ in diatoms are still insufficiently explored.

Recent advances in structural biology and sequencing technologies have provided photosynthetic genomes and protein landscapes (Armbrust et al. 2004; Bowler et al. 2008). These findings reveal that, during the evolution of diverse photosynthetic organisms from cyanobacteria through primary and serial endosymbiosis, the photosystems (PS) remain highly conserved, whereas diverse LHCs have evolved (Croce and Van Amerongen. 2020; Iwai et al. 2024). Diatoms display an expanded repertoire of LHCs compared to green lineages, which is reflected in the genomic of LHC-encoding genes and in the number of LHCs in PSII–LHCII and PSI–LHCI supercomplexes (Nagao et al. 2019, 2020, 2022; Pi et al. 2019; Wang et al. 2019; Xu et al. 2020; Feng et al. 2023, 2025; Zhao et al. 2023; Kato et al. 2024). In the genome of the diatom *Chaetoceros gracilis* (*C. gracilis*), 46 LHCs have been identified, along with a red-lineage chlorophyll *a*/*b*-binding-like protein (RedCAP), a unique member of the LHC superfamily. Phylogenetic analysis revealed that these LHCs can be categorized into six subfamilies: Lhcr, Lhcz, Lhcq, Lhcf, Lhcx, and homologs of CgLhcr9 (Kumazawa et al. 2022). Of the LHCs in the *C. gracilis* genome, only 26 have been found to associate with PSI/II supercomplexes. The remaining 20 LHCs are either not expressed or localized in a peripheral LHC pool. Lhcx proteins responsible for NPQ have not been found in the supercomplex structures to date (Grouneva et al. 2011; Giovagnetti et al. 2022). Lhcx6_1 was recently found tightly bound to PSII in the diatoms *Thalassiosira pseudonana* (*T. pseudonana*) (Feng et al. 2023) and *C. meneghiniana* (Zhao et al. 2023). Its phylogenetic analysis indicates that Lhcx6_1 is distantly related to canonical Lhcx1 and that its absence does not impair NPQ induction, suggesting that Lhcx6_1 is more likely involved in light harvesting rather than photoprotection (Nakayasu et al. 2024).

Building on our previous extensive characterization of the diatom *C. gracilis*—including genome and transcriptome assembly and structural analysis of both photosystems—, we used the recently developed CRISPR/Cas9 technology to characterize the functions of the LHC gene that are not observed in the current cryo-electron microscopy (cryo-EM) structures. We found that deletion of Lhcf2, a protein localized in the peripheral PSII antenna, abolished qE-NPQ. This finding is surprising, demonstrating the critical role of a non-Lhcx-like LHC protein in diatom photoprotection.

## Results

### Lhcf2 constitutes a major component of the peripheral light-harvesting antenna for PSII

The *C. gracilis* genome encodes 14 Lhcf proteins, among which six (Lhcf1, 4, 5, 6, 7, and 13) have been identified within the PSII–LHCII Core_2_S-tetramers_2_M-tetramers_2_ (C_2_S_2_M_2_) supercomplex, constituting the majority of the LHCII subunits (Nagao et al. 2019, 2022; Pi et al. 2019). Lhcf3 is localized to the outermost layer of the PSI–LHCI supercomplex, as revealed by the recent cryo-EM structural analyses (Xu et al. 2020). The status of the remaining seven Lhcf proteins (Lhcf2, 8, 9, 10, 11, 12, and 14) was not resolved by cryo-EM, suggesting that they are either loosely associated with the PSII–LHCII C_2_S_2_M_2_ supercomplexes as PSII–LHCII C_2_S_2_M_2_L_n_ (PSII–LHCII with Loosely-bound LHCII) supercomplexes or expressed at low abundance (Supplementary Fig. 1). Remarkably, Lhcf2 and Lhcf12 are abundantly located in the LHC dimer band as shown by clear native polyacrylamide gel electrophoresis (CN-PAGE) (Zhou et al. 2024). *Lhcf12* has four gene copies (*Lhcf12a–d*), (Supplementary Fig. 2). Guide RNAs (gRNAs) were designed to target conserved regions within the *Lhcf2* gene coding sequence, and five independent knockout mutants (*lhcf2*) were successfully generated using an established transformation protocol (Fig. 1a, Supplementary Fig. 3) (Ifuku et al. 2015; Kumazawa et al. 2025). *lhcf2* mutants demonstrate pale brown coloration but grow in a photoautotrophic medium (Fig. 1b).

**Fig. 1.**
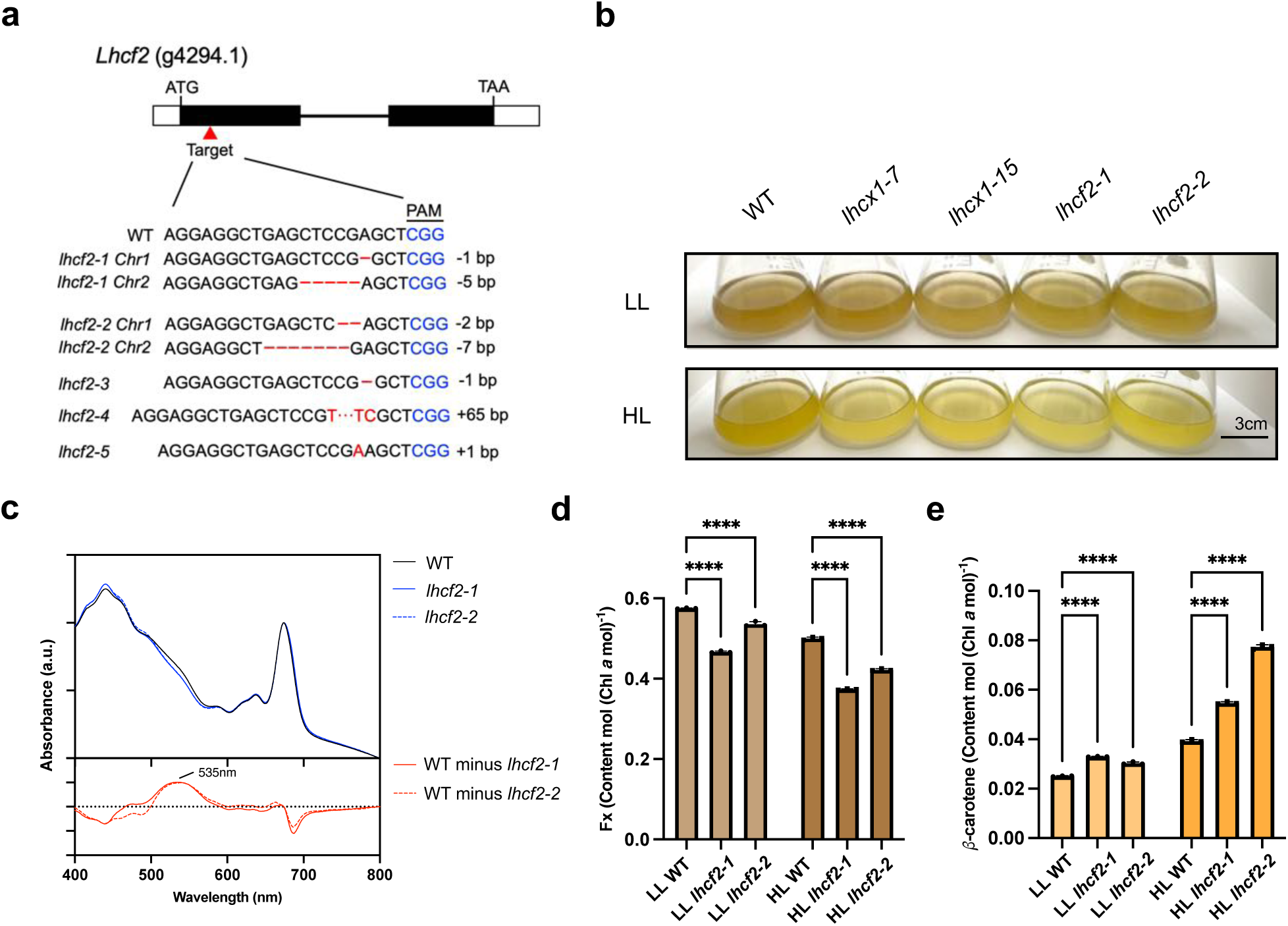
Integrated analysis of the LHC depletion phenotype in *lhcf2* mutants. **a)** Targeted mutagenesis of the *Lhcf2* gene locus was achieved using the CRISPR/Cas9 system. An sgRNA was designed to target a site within the first exon, successfully disrupting gene function. Exons are depicted as black boxes, introns as horizontal lines, and untranslated regions as white boxes. **b)** Comparative photographs of WT and *lhcf2* mutants grown under LL and HL conditions. **c**) room-temperature absorption spectrum (above) and differential spectrum (below) between WT and *lhcf2* mutants. All spectra were rigorously normalized to the absorption at the maximum of the Qy peak. **d-e**) pigment content of Fx (d) and *β*-carotene (e), respectively, relative to Chl *a* quantified by HPLC. Data represent mean ± SD from *n* = 3–4 biological replicates. Statistical significance was determined using two-way ANOVA followed by Dunnett’s multiple comparisons test (*p* < 0.0001, indicated by four asterisks: ****)

To assess the effect of Lhcf2 deletion on the light-harvesting antenna, room-temperature absorption spectra of the wild-type (WT) and two *lhcf2* mutants were analyzed (Fig. 1c). All strains displayed characteristic *C. gracilis* spectral features, including major chlorophyll *a* (Chl *a*; ∼674 nm), minor chlorophyll *c* (Chl *c*; ∼635 nm), and broad spectral shoulders attributable to diadinoxanthin (Ddx; 450–500 nm) and fucoxanthin (Fx; 500–570 nm) (Nagao et al. 2007, 2013b, 2018; Akimoto et al. 2014). Remarkably, a marked reduction in the absorbance shoulder around 535 nm was observed in both two *lhcf2* mutants compared to the WT. This blue–green region corresponds to the absorption spectrum of Fx, a carotenoid pigment that is uniquely associated with diatom LHCs (Nagao et al. 2013b). HPLC analysis shows that the Fx content in *lhcf2* mutants was consistently lower relative to WT under both light conditions (Fig. 1d, Supplementary Fig. 4). In contrast, β-carotene, was markedly elevated in *lhcf2* mutants, under HL (Fig. 1e) and is a biosynthetic precursor of Fx in diatoms (Bai et al. 2022). Taken together, Lhcf2 deletion reduces the cellular Fx pool as a consequence of diminished LHC abundance.

Changes in light-harvesting antenna were precisely evaluated using 77 K fluorescence emission spectroscopy (Elrad et al. 2002). The fluorescence emission ratio at 687 nm to 710 nm (F687/F710) indicates the relative contribution of PSII-associated antenna chlorophylls compared to PSI (Nagao et al. 2018). In *lhcf2* mutants, this ratio was significantly lower than in the WT (Fig. 2a), indicating either a decreased PSII antenna system size or an altered PSII/PSI ratio. Chlorophyll fluorescence induction was monitored in the presence of DCMU (Fig. 2b–d). Under LL conditions, the functional antenna size was reduced by 12.3 ± 1.0 % in *lhcf2-1* mutant and 17.9 ± 3.0 % in *lhcf2-2* mutant compared with WT. In WT grown under HL conditions, the PSII functional antenna size decreased by 7.4 ± 3.1 % relative to LL-grown cells, as a typical photo-acclimation response in diatoms. Remarkably, under HL, the functional PSII antenna size was drastically reduced by 19.6 ± 2.2 % in the *lhcf2-1* mutant and 22.3 ± 1.2 % in the *lhcf2-2* mutant compared with WT under HL conditions; thus, Lhcf2 is critical for the structural PSII functional antenna size.

**Fig. 2.**
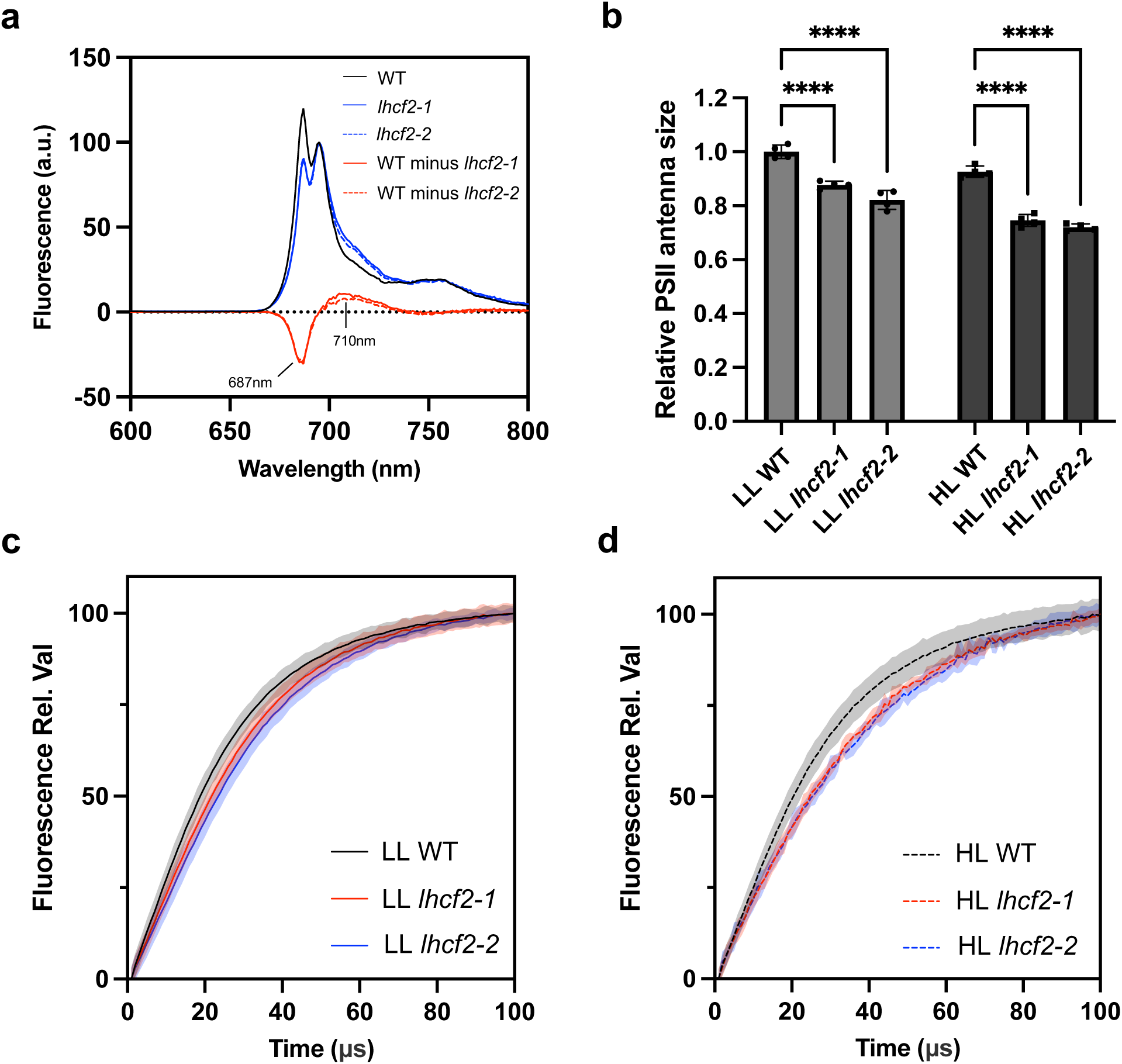
Reduced PSII functional antenna size in the *lhcf2* mutants. **a)** 77K fluorescence emission spectrum and their differential spectrum between WT and *lhcf2* mutants. **b-d**) Chl fluorescence. The reciprocal of the time to reach two-thirds of the total fluorescence rise (T_2/3_), as induced by a short (100 μs) as a quantitative measure of the PSII functional antenna size (b). Representative variable Chl fluorescence induction traces for WT and mutants are shown in c and d. Data represent mean ± SD from *n* = 3–4. Statistical significance was determined by two-way ANOVA followed by Dunnett’s multiple comparisons test (*p* < 0.0001, ****).

### Loss of Lhcf2 causes Lhcx1 reduction in *lhcf2* mutants

CN-PAGE is widely used in photosynthetic research to separate protein complexes while preserving their activity and native structure (Kameo et al. 2021; Zhou et al. 2024). To evaluate the effects of Lhcf2 deletion, deoxycholate (DOC)-based CN-PAGE was performed on thylakoid membranes isolated from WT and *lhcf2* mutants. In WT, thylakoid was separated into distinct bands, from highest to lowest molecular mass: PSI–LHCI and PSII–LHCII supercomplexes, followed by LHC trimer, dimer, monomer, and free pigments (Fig. 3a). Band identities were assigned based on two-dimensional (2D)-PAGE (Fig. 6a, Supplementary Fig. 5a–b) and reported data (Nagao et al. 2013a; Zhou et al. 2024). Notably, the LHC tetramers were not dissociated from the C_2_S_2_M_2_ supercomplex since a relatively low concentration of mild detergent (α-DDM) was used in our CN-PAGE analysis. A marked reduction was observed in the LHC dimer and monomer bands in the *lhcf2* mutant, and the LHC trimer band was almost completely absent. The apparent molecular mass of the isolated PSII–LHCII C_2_S_2_M_2_ supercomplex remained unchanged in the *lhcf2* mutant, suggesting that the oligomeric LHC complex containing Lhcf2 corresponds to the loosely associated peripheral antenna components of PSII.

**Fig. 3.**
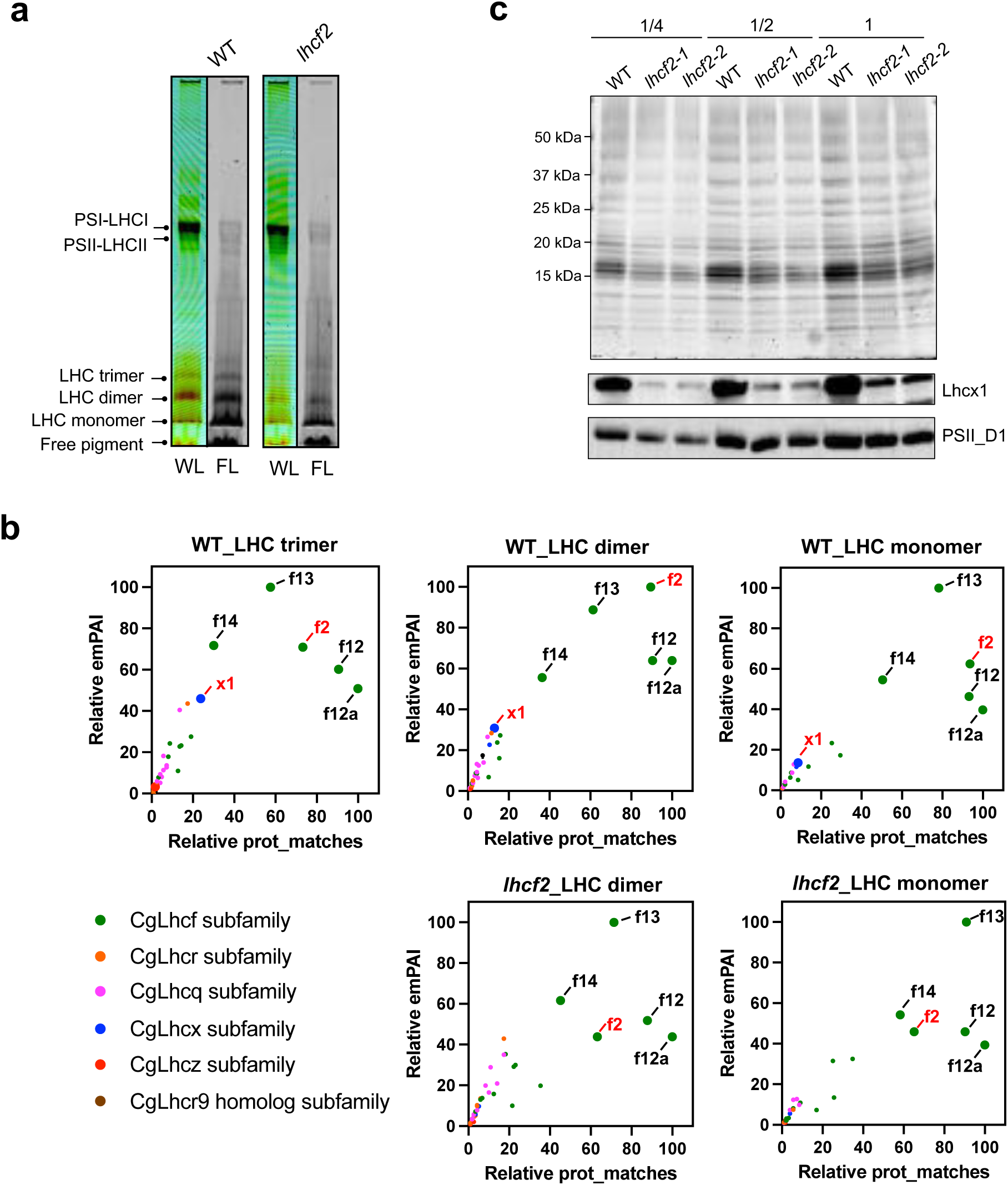
Recognition of Lhcx1 and other LHC components in WT and *lhcf2* mutants. **a)** CN-PAGE analysis (4-13% gradient gel) comparing WT and *lhcf2* mutants. Major LHC oligomeric states (monomers to trimers) and PSI–LHCI/PSII–LHCII supercomplexes are indicated. White-light) and fluorescence (FL) images are presented. **b**) MS analysis of the CN-PAGE gel between WT and *lhcf2* mutants. **c**) SDS-PAGE and immunoblotting analysis against Lhcx1 and the PSII D1 subunit.

We conducted mass spectrometry (MS) on the LHC monomer, dimer, and trimer bands from WT, LHC monomer and dimer bands in the *lhcf2* mutant. The LHC trimer band was almost absent in the *lhcf2* mutant and was therefore not analyzed. As each band was analyzed from a single gel fragment, the exponentially modified protein abundance index (emPAI) values were used to infer the relative abundance of each protein within a given band. However, owing to the high sequence similarity among several LHC isoforms—particularly among Lhcf2, Lhcf12 (originated from *Lhcf12b*/*c*/*d* gene), and Lhcf12a (originated from *Lhcf12a* gene) (Supplementary Fig. 6)—another MS parameter, prot_match (the number of peptides matched to a specific protein), was also considered. By combining both of the scores, semi-quantitative (emPAI) and qualitative (prot_match) assessments of the LHC composition were performed. The results showed that, in LHC trimer, dimer, and monomer bands of WT, Lhcf2 is identified and other LHC components such as Lhcf12, Lhcf12a, Lhcf13, and Lhcf14 are also identified and hold high relative emPAI (Fig. 3b). Noticeably, Lhcx1 is also identified in these three bands, with higher relative emPAI in LHC trimer band but lower relative emPAI in the LHC dimer and monomer bands.

Although the relative emPAI scores for Lhcf2 decreased by 56% in dimer band and 26% in monomer band compared to WT, the presence of Lhcf2 was still indicated (Fig. 3b). However, specific knockout of Lhcf2 was rationalized by further examination of identified peptide sequences: Tryptic digestion of Lhcf2 results in 18 peptides (Supplementary Fig. 7a). Detailed sequence comparison revealed only three specific peptides (AGVHLPGTIDK, DVTGEAEFPGDFR, and IAQLAFLGNIITRAGVHLPGTIDK), whereas the other peptides were conserved among other LHC isoforms. In the WT trimer band, 13 Lhcf2 peptides were identified, of which nine peptides were shared with other LHCs, and two were unique to Lhcf2 with PSM (peptide-spectrum matches) counts of 4 and 6, respectively (Supplementary Fig. 8a–b). The WT dimer and monomer band consistently yielded 14 Lhcf2 peptides each, with all three unique (dimer spectral counts: 8, 31, and 1; monomer spectral counts: 12, 35, and 2). The Lhcf2 is localized across the LHC trimer, dimer, and monomer fractions in WT. In contrast, in the LHC dimer band of *lhcf2* mutant, nine peptides corresponding to Lhcf2 were detected, all of which were conserved among other LHC proteins, with no unique peptides observed. In the *lhcf2* mutant monomer band, 12 peptides corresponding to Lhcf2 were identified, of which 11 were shared with other LHC proteins, and only one unique peptide, “DVTGEAEFPGDFR,” was detected with only a single spectral count, which is likely owing to sample contamination and far fewer than the 35 counts observed in WT. These confirm that the LHC trimer, dimer, and monomer, which are normally enriched with Lhcf2 in WT, contain no detectable Lhcf2 in the *lhcf2* mutant, confirming the success of the knockout.

Intriguingly, Lhcx1 is substantially reduced in the *lhcf2* mutant. Lhcx1 tryptic digestion is predicted to generate 19 peptides (Supplementary Fig. 7b), four (EEYAPGDLR, AQEGWVDPADCPVDQPGLLK, FDPFGLMPEDPEEFDIMQTK AQEGWVDPADCPVDQPGLLKEEYAPGDLR) of which are unique to Lhcx1. In WT trimer band, Lhcx1 was highly represented: 10 peptides, including all four unique to Lhcx1 with PSM counts of 3, 2, 5 and 2, respectively (Supplementary Fig. 8c–d). In the WT dimer band, 10 peptides corresponding to Lhcx1 were detected, including four unique peptides with spectral counts of 3, 2, 3, and 1. In the WT monomer band, nine peptides corresponding to Lhcx1 were identified, including three unique peptides with spectral counts of 2, 2, and 1. In contrast, in the LHC dimer and monomer band of the *lhcf2* mutant, five peptides corresponding to Lhcx1 were detected, and only one unique peptide, “EEYAPGDLR” which was detected with only a single spectral count likely owing to sample contamination, far fewer than that observed in WT. These MS data above indicate that Lhcx1 decreased in the *lhcf2* mutant. Consistently, SDS-PAGE followed by immunoblotting using an anti–Lhcx1 antibody revealed a substantial decrease in Lhcx1 protein levels in *lhcf2* mutants compared to WT (Fig. 3c).

We confirmed that the *Lhcx1* gene sequences in the *lhcf2* mutants are essentially identical to that of the WT, indicating that the *Lhcx1* gene in *lhcf2* mutants was not affected by any off-target or insertion effects of the CRISPR/Cas9 system (Supplementary Fig. 9a–b). Furthermore, the transcript levels of the *Lhcx1* gene in *lhcf2-1* and *lhcf2-2* mutants were comparable to those in the WT (Supplementary Fig. 9c). These data suggest that Lhcf2 is required for the accumulation of Lhcx1.

### Lhcf2 is essential for nonphotochemical quenching function

Lhcx1 is essential for NPQ in *C. gracils* since its knockout almost abolishes NPQ induction under LL and HL-grown conditions (Kumazawa et al. 2025). The reduction of Lhcx1 in *lhcf2* mutants suggests that NPQ should be affected in *lhcf2* mutants. Analysis of maximal NPQ revealed that all five *lhcf2* mutants exhibited markedly reduced NPQ under blue actinic light compared to WT, consistent with observations in *lhcx1* mutants (Fig. 4a). NPQ was subsequently characterized by monitoring time-dependent NPQ induction followed by dark recovery (induction curve) and light intensities response of NPQ (light curve). In the induction curve of WT, NPQ increased rapidly by blue actinic light illumination, reaching a maximum of ∼1.4 (Fig. 4b) and ∼2.4 (Fig. 4c) within 2 minutes of actinic light exposure in cells grown under LL and HL conditions, respectively. In contrast, *lhcx1* and *lhcf2* mutants exhibited minimal NPQ, reaching only ∼0.1–0.2 even at the end of the actinic light treatment. Moreover, the majority (>90%) of the NPQ developed in WT was relaxed during the dark recovery phase, whereas <50% of NPQ was reversible in the *lhcx1* and *lhcf2* mutants. The NPQ was further tested under varying light intensities and the *lhcx1* as well as *lhcf2* mutants consistently showed substantially reduced NPQ compared to the WT (Fig. 4d–e). Similar results were also observed under red actinic light (Supplementary Fig. 10). These findings indicated that the *lhcx1* and *lhcf2* mutants have largely lost the capacity to induce qE-type NPQ. The small amount of NPQ observed under actinic light in the mutants is likely attributable to qI rather than qE, as evidenced by the slower relaxation kinetics during the dark recovery phase relative to the WT (Nawrocki et al. 2021).

**Fig. 4.**
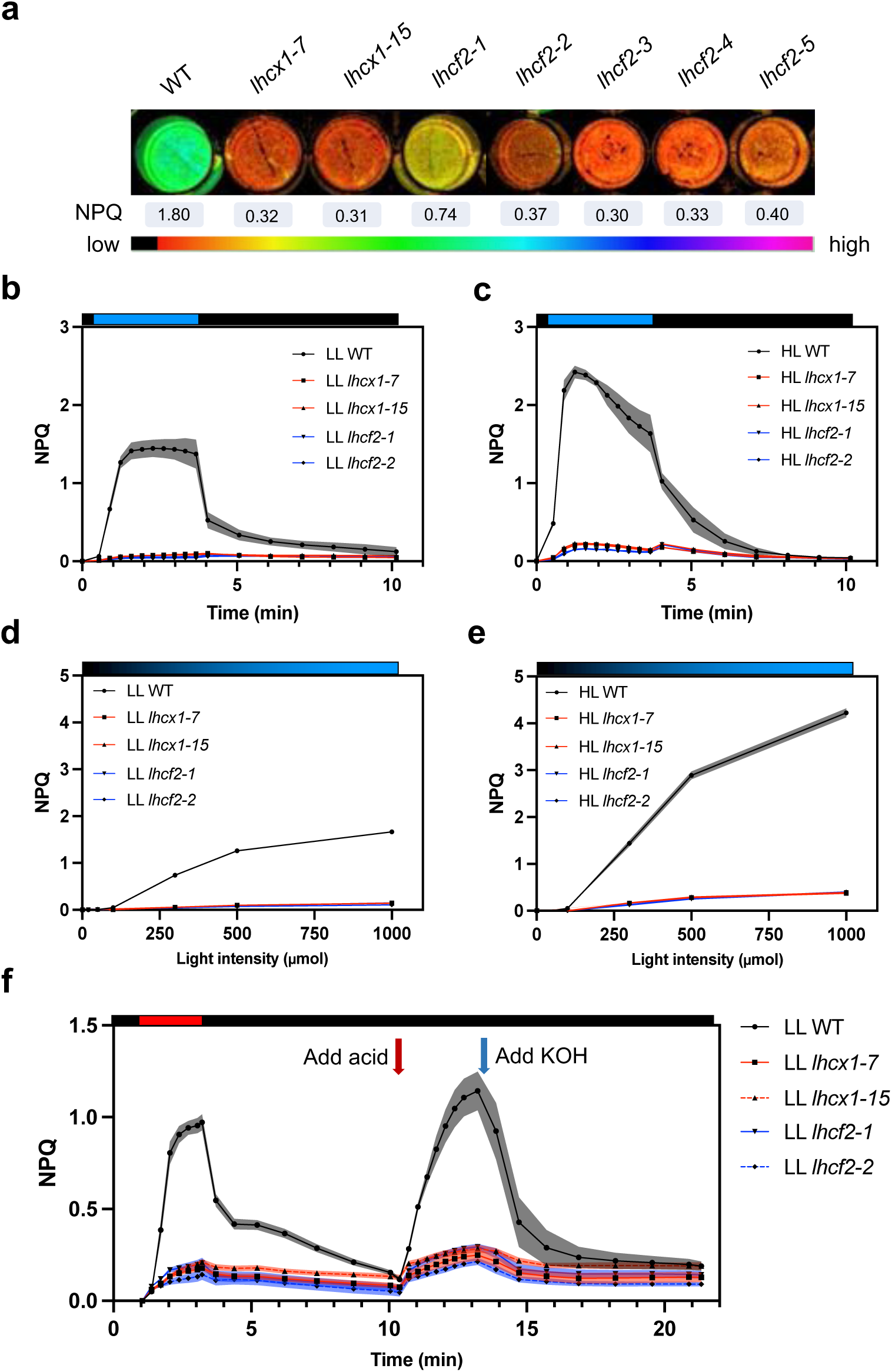
Assessment of light-induced, lumen acidification-induced NPQ response in WT and mutants. **a)** Maximum of NPQ under blue actinic light for WT and mutants grown under LL condition. **b, d**) Induction curve (b) and light curve (d) of NPQ measured by blue actinic light among WT, *Lhcx1,* and *Lhcf2* mutants grown under LL conditions. **c, e**) Induction curve (c) and light curve (e) of NPQ measured by blue actinic light among WT, *lhcx1,* and *lhcf2* mutants grown under HL condition. **e**) lumen acidification-induced NPQ response in WT and mutants. NPQ measurements in WT, *lhcx1,* and *lhcf2* mutants following artificial acidification of the thylakoid lumen in darkness. Cells grown under LL conditions were dark-adapted and subjected to weak red light (indicated by the black bar), during which acetic acid (final concentration 5 mM) was added at 10 min to induce lumen acidification, followed by neutralization with KOH (5 mM final concentration) at 13 minutes. NPQ was continuously monitored using a dual-PAM fluorometer. Data represent mean ± SD from *n* = 3–4 biological replicates.

The molecular mechanisms underlying NPQ have been extensively characterized in land plants and green algae. In land plants, NPQ is PsbS–dependent, wherein the PsbS protein acts as a ΔpH sensor, triggering conformational changes that lead to the clustering of LHCII and the formation of energy-dissipating states upon lumen acidification (Ruban and Wilson. 2021). This process involves the xanthophyll cycle, particularly the conversion of violaxanthin to zeaxanthin, which enhances thermal energy dissipation. In contrast, green algae utilize an Lhcsr–dependent NPQ mechanism, in which Lhcsr proteins bind pigments and serve as pH sensors and quenchers, though PsbS of green algae has its ΔpH-sensing ability (Dewa et al. 2025). Upon lumen acidification, Lhcsr undergoes conformational changes that facilitate energy dissipation independent of LHCII aggregation. These distinctions have been further supported by experimental evidences: in plants, enhancing the pH gradient across the thylakoid membrane can recover NPQ even in the absence of PsbS (Johnson and Ruban. 2011), whereas in green algae, artificially acidifying the lumen in the dark can fully restore NPQ in WT, but fails to do so in Lhcsr3–deficient mutant (Liu et al. 2024).

To determine whether diatoms adopt a similar mechanism, we manipulated lumen acidification and recovery experiments under dark conditions on WT, *lhcx1* and *lhcf2* mutants, using acetic acid and KOH, as described in previous studies (Tian et al. 2019; Liu et al. 2024). In WT, the NPQ value reached 0.98 at the end of actinic light exposure but increased further to 1.16 during artificial lumen acidification (Fig. 4f). This strong NPQ induction in the dark is reminiscent of the response observed in green algae (Liu et al. 2024). However, in the *lhcx1* mutant where Lhcx1 is absent but Lhcf2 is still present, NPQ remained consistently low (∼0.2), regardless of actinic light or artificial lumen acidification, mirroring the phenotype of Lhcsr3–deficient mutants in green algae. Similarly, in the *lhcf2* mutants which lack Lhcx1 and Lhcf2, NPQ could not be restored, showing a response nearly identical to that of the *lhcx1* mutants. Collectively, these results suggest that diatoms, like green algae, rely on a pre-existing quencher likely involving Lhcx1 that is rapidly activated by low lumen pH.

Since the thermal energy dissipation shortens the lifetime of singlet-excited chlorophyll and reduces the amount of excitation energy transferred to the reaction centers under HL conditions, the *lhcf2* mutants are more susceptible to photoinhibition than the WT (Elrad et al. 2002). To assess this outcome, Fv/Fm (the maximum quantum yield of PSII) was measured before and after one hour of HL exposure (∼200 μmol photons m^-2^ s^-1^) in the presence of lincomycin to suppress PSII repair. The *lhcf2* mutants exhibited a more rapid decline in PSII activity than the WT, with half-times of 38.8 minutes (*lhcf2-1*) and 38.0 minutes (*lhcf2-2*), than the 46.8 minutes observed in WT cells (Supplementary Fig. 11a), indicating enhanced photoinhibition in the absence of Lhcf2. Despite the reduced light-harvesting antenna, *lhcf2* mutants exhibited remarkable phenotypic resilience, maintaining a growth rate comparable to that of the WT under LL and HL conditions (Supplementary Fig. 11b-c).

### The trans-thylakoid pH gradient (ΔpH) and xanthophyll cycle operate normally in *lhcf2* mutants

The major component of NPQ in diatoms, qE, requires the concerted actions of ΔpH, de-epoxidized xanthophyll carotenoids, and Lhcx proteins. To elucidate the reason for the failure of NPQ in *lhcf2* mutants, we systematically investigated these prerequisites. ΔpH was assessed using the probe 9-aminoacridine (9-AA) (Seydoux et al. 2022). A decrease in thylakoid lumen pH induces structural changes in lumen-localized 9-AA molecules, shifting their fluorescence emission wavelength outside the detection range of the fluorescence detector and resulting in fluorescence quenching from the perspective of the detector, and thus a reduction in 9-AA fluorescence under actinic light serves as an indirect measure of ΔpH across the thylakoid membrane (Schuldiner et al. 1972; Schreiber and Klughammer. 2009). By employing the sophisticated Dual-PAM system, which uniquely enables the simultaneous monitoring of 9-AA and chlorophyll fluorescence signals, NPQ and ΔpH formation were tracked in real time. In WT cells grown under LL conditions, chlorophyll fluorescence exhibited a sharply increase upon actinic light exposure, followed by a rapid decline (Supplementary Fig. 12a). After ∼20 seconds, chlorophyll fluorescence stabilized and then gradually declined until the end of illumination. 9-AA fluorescence displayed a rapid quenching immediately after light exposure, reaching a minimum value at ∼20 seconds, indicating the rapid establishment of trans-thylakoid ΔpH, and this signal subsequently recovers (Fig. 5a, Supplementary Fig. 12a). The *lhcx1* and *lhcf2* mutants grown under LL conditions displayed similar 9-AA fluorescence kinetics pattern; however, the extent of 9-AA fluorescence quenching was greater in *lhcx1* and *lhcf2* mutants than that in WT, suggesting that the mutants were capable of establishing an enhanced ΔpH than WT (Fig. 5a). Despite that when cells were grown under HL conditions, the magnitude of the ΔpH was comparable among WT, *lhcx1,* and *lhcf2* mutants (Fig. 5b, Supplementary Fig. 12b), these findings rule out defects in ΔpH formation as the cause of NPQ loss in the *lhcf2* mutants.

**Fig. 5.**
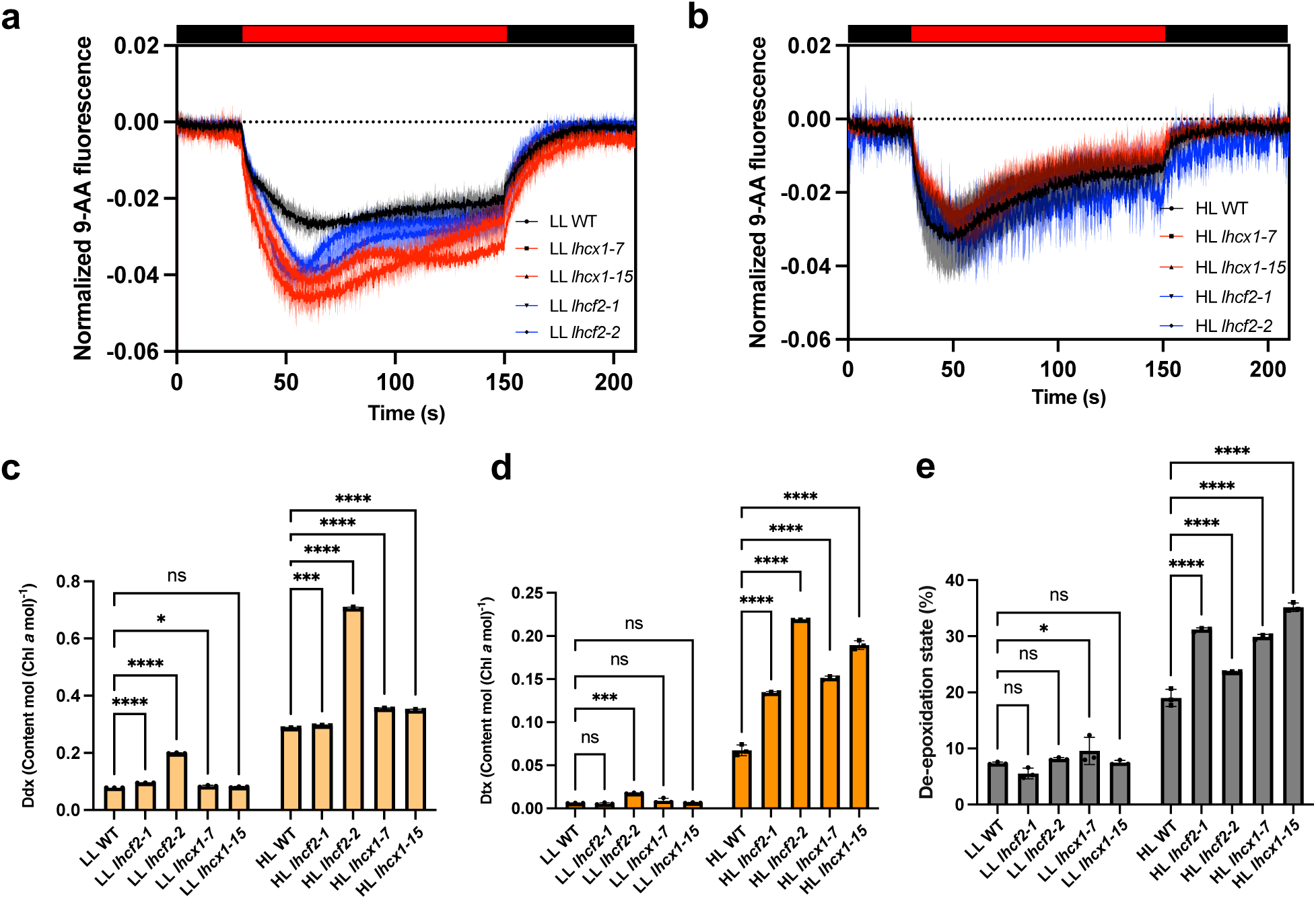
Characterization of NPQ-related components in WT and mutants. **a**, **b**) 9-AA fluorescence quenching kinetics under actinic light measured in WT, *lhcx1,* and *lhcf2* mutants grown under LL (a) and HL (b) conditions using the dual-PAM system. A drop in 9-AA fluorescence indicates formation of a trans-thylakoid pH gradient. **c–e**), relative diadinoxanthin (Ddx), diatoxanthin (Dtx), and de-epoxidation state (DES) of xanthophyll cycle pigments under LL and HL conditions in WT, *lhcx1*, and *lhcf2* mutants. Data in each figure represent mean ± SD with biological replicates of n = 3. Statistical significance was determined by two-way ANOVA followed by Dunnett’s multiple comparisons test (ns, not significant; *p* < 0.001, ***; *p* < 0.0001, ****)

Ddx, Dtx, and the de-epoxidation state (DES) of xanthophyll were analyzed by HPLC (Fig. 5c–e). Under LL conditions, the DES value in *lhcx1* and *lhcf2* mutant were comparable to that of the WT, whereas *lhcf2-2* displayed an even higher DES than the WT (Fig. 5e). HL-grown WT cells showed a substantially elevated DES (∼18%) relative to LL-grown cells (∼7%), correlating with NPQ capacity (Fig. 4b–e). *lhcf2* mutants grown under HL exhibited even higher DES levels than the WT (Fig. 5e), which is consistent with *lhcx1* mutants (Kumazawa et al. 2025), confirming that the xanthophyll cycle remained functional and was not responsible for the loss of NPQ in *lhcf2* mutants. Lhcx proteins have been proposed to act as major reservoirs of protein-bound Ddx (Bailleul et al. 2010); thus, loss of Lhcx1 is expected to release Ddx and enhance its conversion to Dtx. Given that Lhcx1 is largely reduced in *lhcf2* mutants, these results supports that the absence of qE in *lhcf2* mutants is not owing to impaired ΔpH formation or xanthophyll cycle activity, but rather the result of Lhcx1 protein loss.

### Lhcx1 and Lhcf2 co-migrate in CN/SDS-PAGE analysis

Since the *Lhcx1* gene sequence and its corresponding transcript levels were found to be unchanged in the *lhcf2* mutants, the loss of Lhcx1 protein is most likely attributable to post-transcriptional events. This strongly suggests that Lhcf2 may function as a chaperone or structural partner required for the stable accumulation of Lhcx1 protein, potentially through direct physical association. MS showed that Lhcf2 and Lhcx1 are present in the LHC monomer, dimer, and trimer fractions on CN-PAGE of the WT thylakoids (Fig. 3b). To further confirm the distribution pattern of Lhcf2 and Lhcx1, we performed immunoblotting using an anti–Lhcx1 antibody following second-dimension SDS-PAGE. The immunoblot signal showed that Lhcx1 migrates at molecular weight of ∼20 kDa, and its position corresponds to the expected positions of the LHC monomer, dimer, and trimer bands as observed in CN-PAGE (Fig. 6a). Notably, while Lhcf2 protein spots were visible by Oriole statning-induced fluorescence on the 2D-PAGE gel, Lhcx1 spot could not be detected—likely owing to its inherent low abundance. To compare the distribution patterns of Lhcf2 and Lhcx1, the 2D-PAGE and immunoblot images were merged (Fig. 6b). The resulting overlay suggests that the distribution pattern of Lhcx1 closely mirrors that of Lhcf2: Lhcx1 signals appeared directly above regions where Lhcf2 was present and were absent from regions lacking Lhcf2. These results suggest that Lhcx1 is present in the LHC trimer, dimer, and monomer bands and shares a similar distribution profile with Lhcf2, implying a biochemical interaction between the two. In the *lhcx1* and *lhcf2* mutants without the Lhcx1 protein, 2D-PAGE still revealed the presence of Lhcf2/12 or only Lhcf12 signals (Supplementary Fig. 5a-b). Furthermore, the LHC trimer, dimer, and monomer bands enriched of Lhcf2 in CN-PAGE remained at similar intensities between WT and *lhcx1* mutant, without an obvious reduction seen in the *lhcf2* mutant (Supplementary Fig. 5c). These observations indicate that the dependency between these two proteins appears to be asymmetric: Lhcx1 requires Lhcf2 for stable accumulation, but not vice versa—Lhcf2 can exist either independently or in association with Lhcx1.

**Fig. 6.**
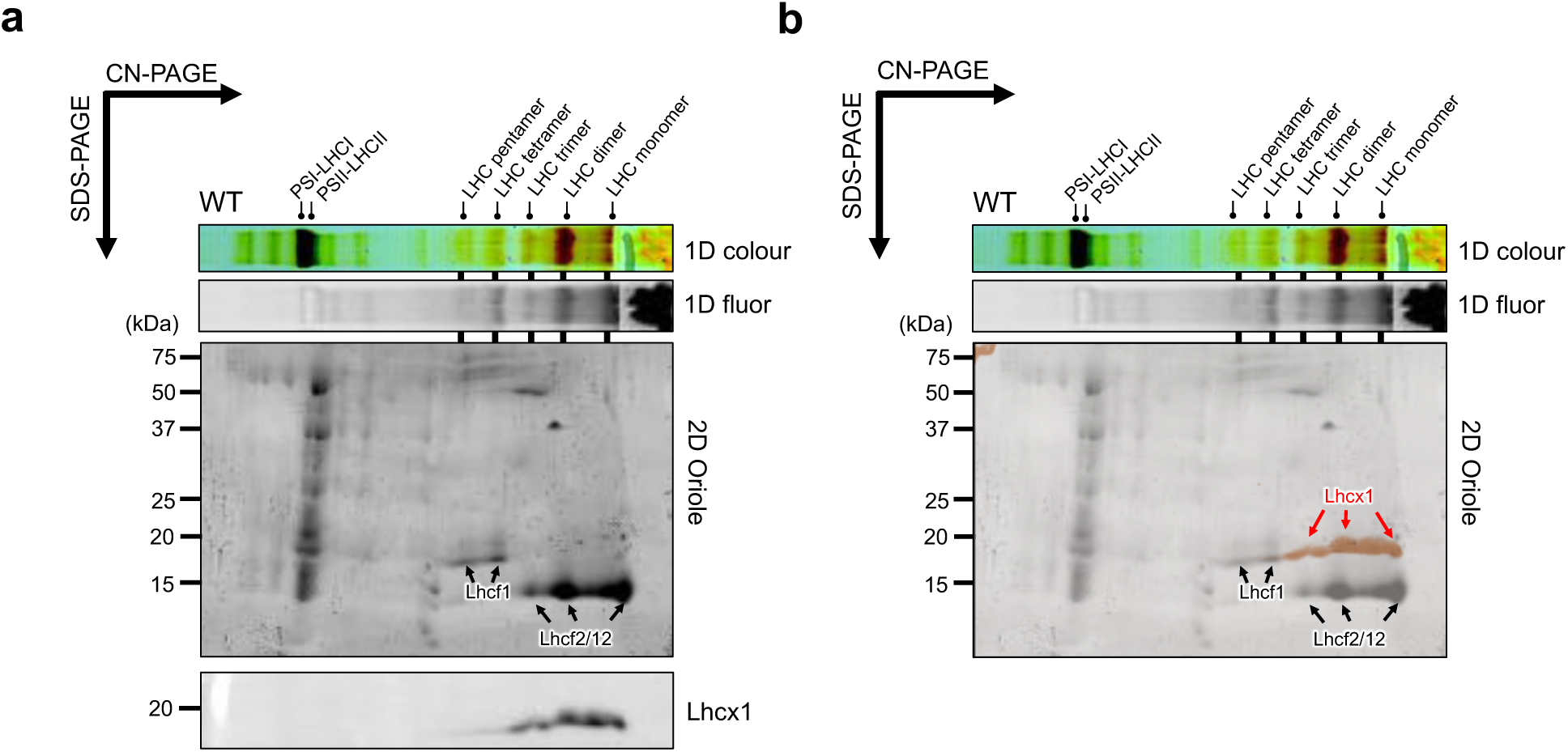
Co-migration analysis reveals Lhcx1 and Lhcf2. **a)** Two-dimensional (2D) CN/SDS-PAGE analysis of WT Oriole staining or immunoblotted with an anti–Lhcx1 antibody. Major LHC with different oligomeric states (monomer to pentamer) and PSI–LHCI/PSII–LHCII supercomplexes are indicated on the CN-PAGE. Lhcf1 and Lhcf2/12 are indicated on the 2D-PAGE. **b**) overlay of oriole-stained gel (black) and anti-Lhcx1 immunoblot (orange). Lhcf1 (black), Lhcf2/12 (black), and Lhcx1 (red) are indicated.

Taken together, these findings support a model in which Lhcx1 is stabilized through incorporation into a specific complex *in vivo*, with Lhcf2 serving as its essential component. In the absence of Lhcf2, Lhcx1 likely fails to incorporate into this stable complex properly and is subsequently degraded, highlighting the broader structural role of Lhcf2 within the LHC protein network. However, it should be noted that this result reflects the oligomeric states of Lhcx1 under *in vitro* biochemical fractionation conditions, which may only partially represent its native states *in vivo*.

### Phylogenetic analysis reveals the evolutionary conservation and relationships of CgLhcf2 and CgLhcx1 across diatom species

In the centric diatom *C. gracilis*, a molecular association between the light-harvesting proteins Lhcf2 and Lhcx1 was identified, potentially forming a functional unit involved in photoprotection. To explore whether this interaction is evolutionarily conserved among other diatoms, we conducted a comprehensive phylogenetic analysis of LHCs based on the open database. For this analysis, we compiled LHC protein sequences from the genomes of several centric diatoms, including *T. pseudonana*, *Thalassiosira oceanica* (*T. oceanica*), and *C. meneghiniana*, as well as from pennate diatoms such as *P. tricornutum*, *Fragilariopsis cylindrus*, *Fistulifera solaris*, and *Pseudo-nitzschia multistriata*. Using these sequences, we inferred a maximum-likelihood phylogenetic tree (Fig. 7). We first categorized the LHC sequences into six major subfamilies based on the classification in *C. gracilis*: Lhcr (orange), Lhcz (red), Lhcf (green), Lhcq (magenta), Lhcx (blue), and CgLhcr9 homologs (brown) (Kumazawa et al. 2022).

**Fig. 7.**
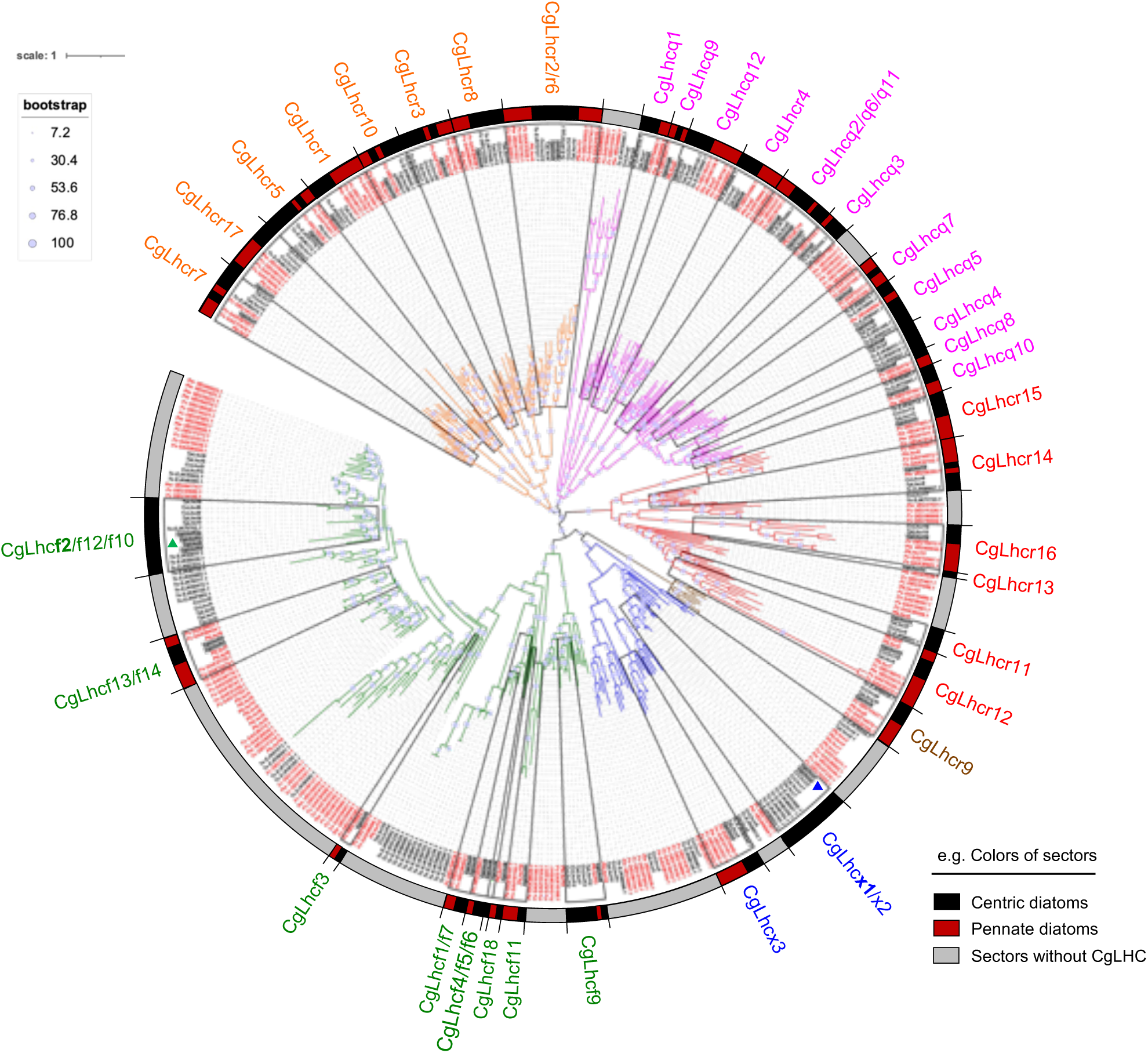
Phylogenetic classification of LHC proteins across centric and pennate diatoms. A maximum-likelihood (ML) phylogenetic tree was constructed using full-length amino acid sequences of LHC proteins derived from *C. gracilis* and seven representative diatom species, including centric diatoms (*Thalassiosira pseudonana*, *Cyclotella meneghiniana*, and *Thalassiosira oceanica*) and pennate diatoms (*Phaeodactylum tricornutum*, *Fragilariopsis cylindrus*, *Fistulifera solaris*, and *Pseudo-nitzschia multistriata*). Based on the *C. gracilis* classification, LHC proteins were grouped into six major families: Lhcr (orange), Lhcz (red), Lhcf (green), Lhcq (magenta), Lhcx (blue), and CgLhcr9 homologs (brown). Subclades were rigorously defined to include at least one *C. gracilis representative*, resulting in 34 annotated subclades labeled with CgLHC. The positions of CgLhcf2 and CgLhcx1 are indicated by green and blue triangles, respectively. Protein sequences originating from centric diatoms are shown in black, those from pennate diatoms in red, and Cg-derived LHCs are marked with gray squares. Colored sectors indicate the taxonomic composition of each clade: black for clades exclusively containing centric diatom sequences, red for pennate-exclusive clades, and gray for clades lacking *C. gracilis* representatives. Node support was estimated by bootstrap analysis and is indicated by circle size.

Within each major group, a further classification was done based on the phylogenetic proximity, ensuring that each subclade included at least one representative LHC from *C. gracilis* (CgLHC) and yielded 34 refined subclades each anchored by a specific CgLHC label that is highlighted on the phylogenetic tree (e.g. CgLhcr7). In each subclade, LHCs from centric diatoms are labelled in black, those from pennate diatoms are in red, and CgLHCs are marked with gray squares. To clearly illustrate the taxonomic origin of each clade, the outermost edge of the tree was annotated with colored sectors: black sectors indicate clades composed of centric diatom LHCs, red sectors indicate those from pennate diatoms, and gray sectors indicate clades that do not contain any CgLHC.

The phylogenetic results revealed that most subclades contained LHCs from centric and pennate diatoms, indicating frequent intermixing of LHC types across these lineages. For instance, in the CgLhcr7 clade, sequences from pennate diatoms such as PtLhcr2, Fs GAX17916.1, Fs GAX24797.1, and Fc OEU14130.1 are present, whereas sequences from To EJK49415.1, TpLhcr19, CcLhcr7, and TpLhcr7 from centric diatoms are also present. This suggests that the majority of CgLHCs are not exclusive to centric diatoms; rather, the presence of closely related LHCs in pennate diatoms appears to be the norm, reflecting a shared ancestral origin despite taxonomic separation. Notably, the presence of a specific CgLHC does not necessarily imply its conservation in other centric diatoms. Specifically, the CgLhcq9 clade includes only LHCs from pennate diatoms, such as Fc OEU16277.1 and PtLhcq3, with no representatives from other centric diatoms. This heterogeneity highlights the complexity of LHC evolution and species-specific adaptations in light harvesting systems.

Of particular note, the subclades containing CgLhcf2 (indicated by the green triangle in Fig. 7) and CgLhcx1 (indicated by the blue triangle in Fig. 7) included LHCs only from centric diatoms. Furthermore, all four centric species surveyed harbor the homologs of CgLhcf2 and CgLhcx1, whereas no such pennate diatom-derived LHCs were identified in those subclades. This stands in stark contrast to most CgLHCs and suggests that Lhcf2 and Lhcx1 are conserved among centric diatoms and lack closely related counterparts in pennate species. Given that protein function is intrinsically tied to the amino acid sequence, the centric diatom-specific nature implies that the molecular interaction between Lhcx1 and Lhcf2 observed in *C. gracilis* is potentially co-evolved and conserved across other centric diatoms.

Based on sequence similarity and phylogenetic proximity, candidate orthologs of CgLhcf2 include TpLhcf3, TpLhcf4, and TpLhcf6 in *T. pseudonana*; To EJK75762.1, To EJK53611.1, To EJK55038.1, and To EJK61531.1 in *T. oceanica;* CcLhcf4 and CcLhcf6 in *C. meneghiniana*. Similarly, candidate orthologs of CgLhcx1 include TpLhcx1 and TpLhcx2 in *T. pseudonana*; To EJK51806.1, To EJK48687.1, To EJK66358.1, To EJK49164.1, To EJK49165.1, and To EJK68853.1 in *T. oceanica*; CcLhcx1 and CcLhcx2 in *C. meneghiniana*. While these assignments are based on robust phylogenetic inference, functional validation through gene-editing approaches such as CRISPR/Cas9 or RNAi will be required to confirm their relationship. Importantly, although no direct orthologs of CgLhcf2 or CgLhcx1 were detected in pennate diatoms, this does not eliminate the possibility that such functionally analogous interactions may occur mediated by different LHCs that are far from CgLhcf2 and CgLhcx1 on the phylogenetic proximity.

## Discussion

We initially identified the essential role of Lhcf2 protein in maintaining NPQ in *C. gracilis*. To further elucidate the cause of NPQ impairment in the *lhcf2* mutant, three major factors in NPQ were systematically examined. After excluding the involvement of ΔpH and xanthophyll cycle, the loss of Lhcx1 was identified as the underlying reason for the abolished NPQ. We therefore sequenced the *Lhcx1* gene in *lhcf2* mutants and confirmed that it was not affected by off-target effects of the CRISPR/Cas9 system. Moreover, the mRNA level of the *Lhcx1* gene in *lhcf2* mutants was comparable to that observed in the WT. These outcomes provide compelling and direct evidence that Lhcf2 plays a crucial role in the regulation of the NPQ key factor, Lhcx1. Additionally, immunoblotting and MS analyses revealed that Lhcf2 and Lhcx1 exhibited highly similar distribution patterns on CN-PAGE, suggesting a potential biochemical interaction between the two proteins. Collectively, these findings support a model wherein the Lhcf2 exists in an oligomeric form assembled together with Lhcx1 and Lhcf2 itself at the periphery of the PSII–LHCII C_2_S_2_M_2_ supercomplex *in vivo*, collectively forming PSII–LHCII C_2_S_2_M_2_L_n_ supercomplex. (Fig. 8).

**Fig. 8.**
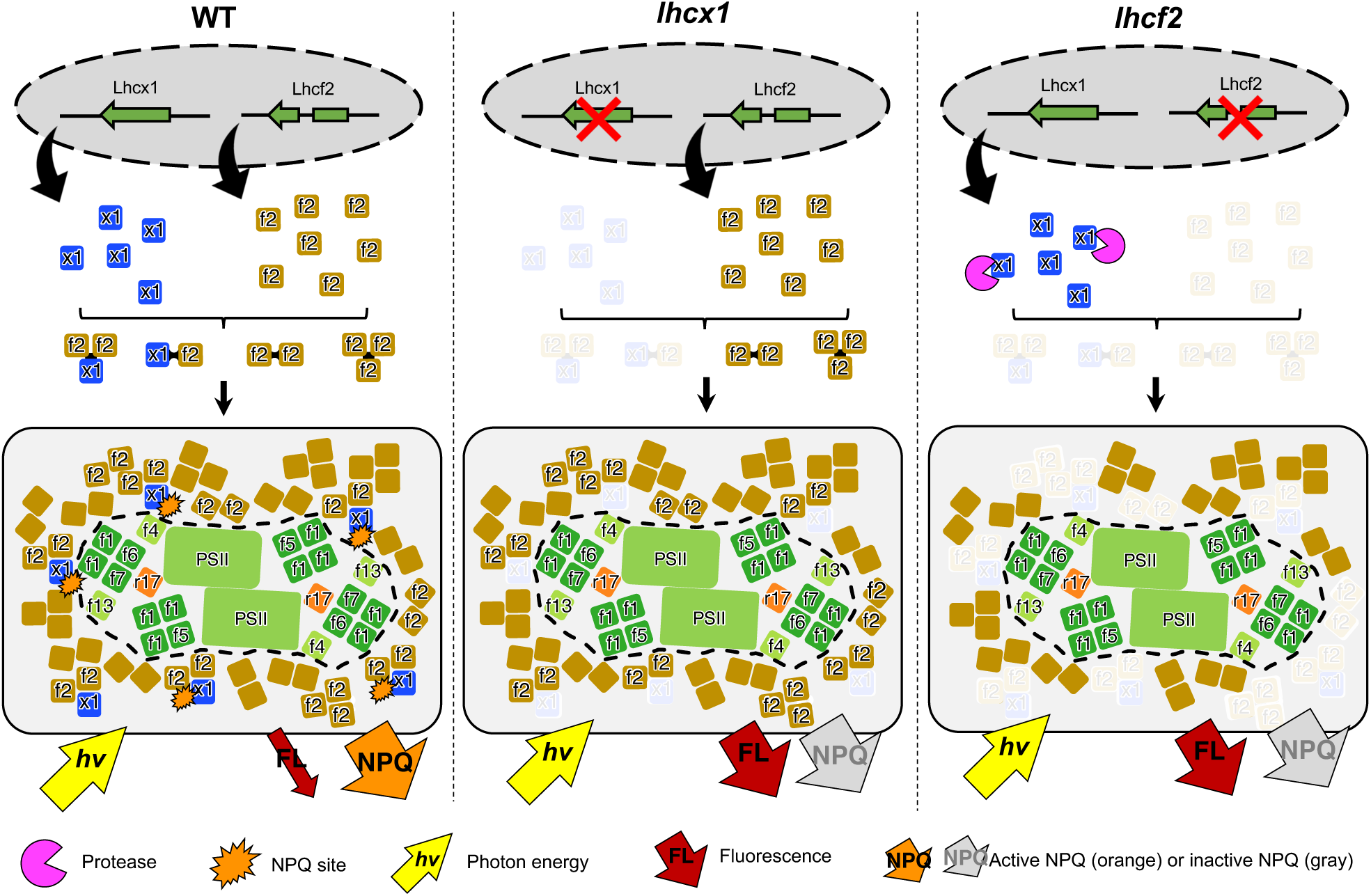
Schematic model detailing PSII–LHCII C_2_S_2_M_2_L_n_ supercomplex organization and the NPQ. In WT, Lhcf2 and Lhcx1 are expressed and likely associate to form diverse peripheral oligomeric complex such as LHC trimer or dimer at the periphery of the PSII–LHCII C_2_S_2_M_2_ supercomplex (indicated by black dashed lines), contributing to energy dissipation via NPQ primarily within the Lhcx1 protein (orange bursts). In the *lhcx1* mutant, Lhcf2 is still present, but the absence of Lhcx1 abolished NPQ and enhances radiative energy loss (red arrows). The *lhcf2* mutants exhibit loss of Lhcf2 which is knocked out and Lhcx1 which is expressed and then degraded by certain protease (magenta sector).

Previous studies in green alga *Chlamydomonas reinhardtii* (*C. reinhardtii*) have demonstrated that the loss of Lhcbm1 abolished qE under HL conditions, despite abundant Lhcsr3 in the mutant, highlighting the indispensable role of certain LHCs in photoprotection (Elrad et al. 2002; Ferrante et al. 2012; Liu et al. 2024). The authors attributed this to the possibility that the absence of Lhcbm1 prevents Lhcsr3 from maintaining a stable quenching conformation or disrupts the interaction between Lhcsr3 and the PSII complex, resulting in reduced energy transfer to the quencher in Lhcsr3 (Liu et al. 2024). However, it is important to note that in *C. reinhardtii*, Lhcbm1 is present in the peripheral antenna and within the S-trimer antenna close to the PSII core (Sheng et al. 2019). Therefore, it remains unclear whether the reduced NPQ observed upon Lhcbm1 loss arises from disruptions within the S-trimer, the peripheral antenna region, or both. Unlike in green algae, structural analyses in the *C. gracilis* show that Lhcf2 is not closely associated with the PSII core at all, such as the S or M-tetramers (Nagao et al. 2022). Instead, all Lhcf2 proteins are loosely associated at the periphery of the PSII-LHCII C_2_M_2_S_2_ supercomplex, nullifying the hypothesis that qE quenching occurs near the PSII core in diatoms and supports the notion that the majority of qE takes place in the peripheral antenna systems far away from PSII (Zhou et al. 2024; Kumazawa et al. 2025). By employing a low concentration of mild detergent and an optimized CN-PAGE system, we demonstrated that a substantial amount of the Lhcx1 protein in higher oligomeric form can be isolated from *C. gracilis* even cultured under LL conditions. However, Lhcx1 is not associated with other LHCs in diatom *P. tricornutum* in biochemical analysis (Grouneva et al. 2011; Giovagnetti et al. 2022). Instead, under LL and intermediate light conditions where Lhcx1 is highly expressed, biochemical approaches such as sucrose density gradient (SDG) centrifugation or CN/BN-PAGE have only detected Lhcx1 in the monomer region (Grouneva et al. 2011; Giovagnetti et al. 2022). Additionally, BN-PAGE analysis of *T. pseudonana* cells grown under LL showed that all Lhcx1 proteins appeared in the free protein band, indicating that they remained in monomeric form as well (Grouneva et al. 2011). However, in cells cultured under HL conditions, BN-PAGE analysis detected Lhcx1 in the free protein band and in the LHC trimer band, as confirmed by MS (Grouneva et al. 2011). Nevertheless, the Mascot score of Lhcx1 in the trimer band was markedly lower than that of other LHC components and substantially lower than its score in the free protein band, suggesting that even under HL conditions, most Lhcx1 remains in monomeric form after biochemical separation (Grouneva et al. 2011). Moreover, a recent study employing CN/SDS-PAGE followed by MS on thylakoids isolated from *T. pseudonana* grown under LL also identified Lhcx1 exclusively in the monomeric form, further supporting the tendency of Lhcx1 to remain monomeric in *T. pseudonana* (Zhou et al. 2024). In fact, although various oligomeric forms of LHCs can be separated from thylakoid membranes using biochemical approaches such as CN/BN-PAGE, ion-exchange chromatography, or SDG centrifugation, the results obtained from these methods can be influenced by isolation method, the type and concentration of detergents used, as well as handling procedures and technical variations, and therefore do not necessarily represent the exact oligomeric state of LHCs in the native LHC pool *in vivo*. Current findings suggest that in *C. gracilis*, Lhcx1 is relatively tightly associated with other LHC proteins. In contrast, the interaction between Lhcx1 and other LHCs might be weaker in *T. pseudonana* and *P. tricornutum*. Given that Lhcx1 is highly conserved between pennate and centric diatoms, the observed differences in how tightly it binds to other LHCs are likely attributable to variations in the amino acid sequences of its LHC partners (Kato et al. 2024). These differences may in turn influence the biochemical separation outcomes. Since Lhcx1 plays an essential role in photoprotection, it is tempting to propose whether the differences in its binding affinity to other LHCs could affect NPQ. If Lhcx1 exists predominantly in a monomeric form, would its NPQ mechanism remain the same? These questions warrant further experimental investigation in the future.

We would like to underscore several caveats in our study and precautionary points that could guide future research directions. First, owing to the limited number of antibiotic types available in our transformation system, we were unable to use an additional antibiotic for constructing complementary strains of the *lhcf2* mutant. Addressing this limitation will be a focus of our future work. Furthermore, plasmids introduced via electroporation could integrate randomly into the genome, potentially disrupting unrelated genes. However, five independent *lhcf2* mutants were obtained, all of which carried frameshift mutations in the Lhcf2 exon and lost NPQ capacity (Fig 4. a–e, Supplementary Fig. 13). These results rule out the possibility that the phenotype was caused by random plasmid insertion or off-target effects of the CRISPR system. Another limitation is the lack of direct evidence demonstrating a physical interaction between Lhcf2 and Lhcx1. While the concomitant loss of Lhcf2 leads to the disappearance of Lhcx1 and both proteins exhibit highly similar distribution patterns on CN-PAGE, these findings merely suggest a high likelihood of biochemical interaction, rather than proving it definitively. The most direct approach would be to visualize a putative Lhcf2–Lhcx1 complex in isolated LHC bands by cryo-EM. However, MS analysis in the *lhcf2* mutant indicates that none of the LHC trimer, dimer, or monomer bands are composed solely of Lhcf2 and Lhcx1; other LHC proteins, such as Lhcf12, Lhcf13, and Lhcf14, are also present. This heterogeneous composition posed a major obstacle for directly resolving the Lhcf2–Lhcx1 interaction via structural analysis. Consequently, future studies employing co-immunoprecipitation (Co-IP) or affinity-tag purification strategies will be necessary to provide direct biochemical evidence for the interaction between Lhcf2 and Lhcx1. Collectively, these observations suggest that Lhcf2 may function as a chaperone-like partner for Lhcx1, facilitating or stabilizing its incorporation within the thylakoid membrane. Once synthesized, Lhcx1 diffuses within the LHC pool of the membrane until it encounters Lhcf2, to which it binds with high specificity. In the absence of a suitable binding site, unbound Lhcx1 is likely unstable and subject to degradation by certain proteases (Fig. 8). Notably, this interaction is highly selective and specific, as even though Lhcf12 shares over 90% amino acid identity with Lhcf2 (Supplementary Fig. 6), its presence in *lhcf2* mutant is not sufficient to compensate for the loss of Lhcf2 as a binding partner of Lhcx1. Based on this information, by comparing the amino acid sequences between Lhcf2 and Lhcf12, it is possible to infer which residues in Lhcf2 are critical for the interaction with Lhcx1. The results indicate that several residues—K94, S113, G114, T118, G130, E138, and P142 in Lhcf2—differ markedly from the corresponding residues in Lhcf12—Y94, P113, Q114, L118, F130, deletion at position 138, and V142. These substitutions represent highly nonconservative changes, suggesting that these amino acids may play crucial roles in the specific interaction between Lhcf2 and Lhcx1. In the future, site-directed mutagenesis could be employed to determine which of these residues are decisive for the interaction. Intriguingly, if the presence of Lhcx1 indeed requires a partner protein for its stabilization, and in the absence of such a partner Lhcx1 fails to accumulate in all diatoms, then all monomeric Lhcx1 proteins isolated through biochemical procedures would essentially be artifact. Considering that monomeric Lhcx1 can be readily detected in diatoms using biochemical methods (Grouneva et al. 2011; Giovagnetti et al. 2022), such findings seemingly contradict the hypothesis mentioned that Lhcx1 cannot exist independently without a stabilizing partner. A plausible explanation for this discrepancy is that Lhcx1 is loosely associated with other LHCs in a subtle manner that allows for stable accumulation, but these interactions are disrupted during biochemical fractionation owing to their labile nature, resulting in the eventual detection of Lhcx1 in monomeric form.

Another open question in diatom photoprotection concerns the structural identity of the NPQ site host. The term “quencher” represents the responsible pigment and the specific protein in which the quencher is located. We focus on the latter. Based on picosecond time-resolved fluorescence analysis, the prevailing model for NPQ in *P. tricornutum* and *C. meneghiniana* proposes the formation of two distinct quenching sites (Q1 and Q2) (Holzwarth et al. 2009; Miloslavina et al. 2009; Chukhutsina et al. 2014), despite their FCP antenna configuration (Goss and Lepetit. 2015). The quenching site 1 (Q1) is localized to detached–antenna complexes largely independent of Dtx. In contrast, quenching site 2 (Q2) is situated in an antenna attached to the PSII and is Dtx accumulation-dependent. Lhcf2 and Lhcx1 are localized in the outer layer of PSII–LHCII C_2_S_2_M_2_L_n_ supercomplex as peripheral antenna; they could theoretically participate in either Q1 or Q2. However, the finding that Lhcx1 and Lhcf2 mutants exhibit an equally abolished NPQ response (Fig. 4a–e) suggests that in *C. gracilis*, Lhcx1 is the indispensable constituent required for the function of Q1 and Q2. The presence of Lhcf2 in the *lhcx1* mutant resulted in comparable residual quenching in *lhcf2* mutants. This indicates that Lhcf2 alone is functionally insufficient to NPQ in the absence of Lhcx1 under conditions of artificial lumen acidification (Fig. 4f), which is consistent with recent spectroscopic characterization (Zheng et al. 2023), and definitively establishes Lhcx1 as the host protein mediating the quenching site. Future elucidation of the *in situ* architecture of the PSII antenna system is needed to structurally verify the configuration of the PSII–LHCII C_2_S_2_M_2_L_n_ supercomplex and directly map the precise localization binding partner of Lhcx1 in *C. gracilis*. More broadly, defining this NPQ mechanism and its significant ecological role in marine resource utilization, as well as its potential application in enhancing the production of valuable metabolites for biotechnology, is necessary.

## Materials and Methods

### Cell culture

The marine centric diatom *Chaetoceros gracilis* (UTEX LB 2658) WT, the *lhcx1* mutants, and the *lhcf2* mutants were grown in a baffled shake flask on a rotary shaker (100 rpm) with ambient air at 20°C under continuous LL illumination (30 μmol photons m^-2^ s^-1^) or HL illumination (∼200 μmol photons m^-2^ s^-1^). Culture was performed in artificial seawater consisting of Daigo’s artificial seawater (Daigo MS, Japan), 1× Daigo IMK medium (Daigo MS, Japan), and 0.2 mmol L⁻¹ Na₂SiO₃. Algae were maintained by weekly or biweekly subculture at a tenfold dilution with fresh growth medium. Cell density was monitored using a Thoma hemocytometer over growth. Cells for experiments were collected during the exponential growth phase.

### CRISPR/Cas9-mediated mutagenesis

CRISPR/Cas9 mutagenesis, including guide RNA design, construction of plasmids expressing Cas9 and guide RNAs, and electroporation-based transformation, was performed using a linearized plasmid construct described in our previous research (Ifuku et al. 2015; Kumazawa et al. 2025). Primers that were annealed for generation of the guide RNA template are provided in Supplementary Data 1. The linearized plasmid contained an Sh ble (*Streptoalloteichus hindustanus* bleomycin resistance) gene expression cassette, which allowed for the selection of successfully transformed clones on IMK agar plates containing 100 µg/mL zeocin (InvivoGen, #ant-an-1, USA). The primary transformants underwent PCR amplification and sequencing of the guide region, yielding either completely WT or mixed sequencing results; note that we designed primers (Supplementary Data 1) from regions without polymorphisms between the two chromosomes to avoid missing the amplification of one allele. Colonies with mixed sequencing outcomes were then cultured in liquid medium and subsequently spread onto selective solid medium for another round of colony formation. The sub-clones derived from these colonies could present with the WT sequence, mixed sequencing results, or unequivocal sequencing results. Therefore, we prioritized the selection of sub-clones that either showed unequivocal sequencing results with the same frameshift mutation on both chromosomes or mixed sequencing results indicating distinct frameshift mutations on each chromosome for further physiological analyses.

### Measurement of photosynthetic physiological parameters

Chlorophyll fluorescence was measured using pulse-amplitude modulation with actinic blue light (455 nm) at an intensity of 300 μmol photons m⁻² s⁻¹. Approximately 4 × 10^6^ cells were harvested and resuspended in 2 ml IMK medium. Measurements were performed using the AquaPen AP-110 fluorometer (PSI, Czech Republic) with the NPQ2 and LC3 protocol, which includes a 200-second light phase followed by a 390-second dark recovery phase. The first saturating pulse of light was applied 10 seconds after the onset of actinic light illumination. Fluorescence was detected through a bandpass filter (667–750 nm). The the intensity of the saturating pulse was set to 30%, corresponding to 900 μmol photons m^-2^ s^-1^. Prior to measurement, cells were dark-adapted for at least 10 minutes. Another type of chlorophyll fluorescence was simultaneously measured using pulse-amplitude modulation with actinic red light (620 nm) at an intensity of ∼1,250 μmol photons m⁻² s⁻¹ in a 10 x 10 mm cuvette using a Dual-PAM-100 (Walz Heinz GmbH, Effeltrich, Germany) with the unit ED-101US/MD (Walz Heinz GmbH, Effeltrich, Germany) for suspensions at room temperature. Before the measurement, the cells were diluted with IMK medium and adjusted to a final concentration of 50 μg/ml Chl *a* + *c* with 1 ml volume.

The Chl fluorescence parameters were calculated as described by Baker (Baker 2008). The maximum quantum efficiency of PSII photochemistry was calculated as Fv/Fm = (Fm – Fo)/Fm, and light-induced regulatory non-photochemical quenching of PSII excitation energy was calculated as NPQ = (Fm – Fm’)/Fm’, where Fo is the minimum fluorescence yield, Fm is the maximum fluorescence yield, and Fm’ is the maximum variable fluorescence yield. Fo and Fm were determined by measuring light (0.1 μmol photons m^−2^ s^−1^) and saturated pulse (20,000 μmol photons m^−2^ s^−1^, 300 ms).

For the analysis of the maximal NPQ, approximately 4 × 10^6^ cells were collected by centrifugation and resuspended in 1 ml IMK medium. The cells were dark-adapted for 10 min prior to the measurements and subsequently exposed to blue actinic light (451 nm, 150 μmol photons m⁻² s⁻¹) for 200-second by using the Hexagon Imaging-PAM (Walz Heinz GmbH, Effeltrich, Germany). The maximal NPQ data were exported from the equipped software as a false-color image. For the measurement of PSII photoinhibition, Fv/Fm was measured by using an AquaPen AP-110 fluorometer system after 20, 40 and 60 min of HL exposure (∼200 μmol·m⁻²·s⁻¹) in the presence of lincomycin (0.5 mg ml^−1^). Before measuring Fv/Fm, the cultures were dark-adapted for 10 min. The measurement of acid-induced NPQ was performed as previously reported (Tian et al. 2019). Acetic acid (1 M) was added to the culture to decrease the pH to 5.5, and 2 M KOH was used to set the pH to 7 (Tian et al. 2019). NPQ was recorded using a DUAL-PAM 100 system with saturating pulses (20,000 μmol photons m^−2^ s^−1^; 300 ms duration) and actinic light (∼1,100 μmol photons m^−2^ s^−1^). Cells were dark-adapted for at least 10 min before measurements. Measurements were repeated on 3–4 biological replicates per genotype.

### Determination of 9-aminoacridine fluorescence

9-aminoacridine (9-AA) fluorescence was measured to determine ΔpH using the DUAL-ENADPH and DUAL-DNADPH units of the DUAL-PAM 100 system (Schreiber U and Klughammer C. 2009; Johnson and Ruban. 2011). Excitation was provided by 365-nm LEDs and fluorescence emission was detected between 420 and 580 nm. Measurements were performed on cells of WT and mutants (5 μg Chl *a* mL^-1^) in the presence of 1 μM 9-AA. Following a 5-min dark period, 9-AA fluorescence quenching was detected during an illumination with actinic light of an intensity of about 1,400 μmol m^−2^ s^−1^ under gentle stirring of the reaction mixture under dark conditions. The extent of 9-AA fluorescence quenching was assessed after correction of the 9-AA signal drift observed. The magnitude of the ΔpH was calculated as ΔF/F value, with F being the 9-AA fluorescence before the onset of actinic illumination and ΔF being the difference between the 9-AA fluorescence before the illumination and the 9-AA fluorescence during the actinic light phase.

### Measurement of functional PSII antenna size

For Chl fluorescence induction analyses, 2-ml cell suspensions were adjusted to a cell density of about 2 × 10^6^ cells/ml. Chlorophyll fluorescence induction curves were measured using a Fluorometer FL 3500 (Photon System Instruments, Drasov, Czech Republic), as described by Kaftan (Kaftan et al. 1999). Following 10 minutes of dark adaptation, chl fluorescence was induced in the presence of 40 μM DCMU using continuous actinic illumination of 100-µsec duration, and Chl fluorescence levels were measured every 1 µsec using a weak pulse-modulated measuring flash. The relative antenna size was determined by calculating the reciprocal of time corresponding to two-thirds of the fluorescence increase (T_2/3_).

### Absorption spectra measured at room temperature

Room-temperature absorption spectra were recorded at 25 °C using a UV-2600 spectrophotometer (Shimadzu, Japan) and normalized to the absorption at the maximum of the Qy peak.

### Steady-State Fluorescence Spectra

Steady-state fluorescence emission spectra were measured at 77K using a spectrofluorometer (FP 6600, JASCO) equipped with a PMU-130 liquid nitrogen cooling unit, with excitation wavelength at 459 nm. The samples were diluted with IMK medium and adjusted to a final concentration of 1 μg/ml Chl a + c. Chlorophyll concentration was determined by extraction with 90% acetone according to the method of Jeffrey (Jeffrey and Humphrey. 1975). Spectra were obtained by integrating five consecutive scans with a sampling interval of 1 nm. All spectra were normalized to the CP47 peak of PSII (693–695 nm), as described by Nagao (Nagao et al. 2010).

### Pigment analyses by high-performance liquid chromatography (HPLC)

Cells were harvested by centrifugation at 1,200 × g for 5 minutes, and the supernatant was discarded. The resulting pellet was immediately frozen in liquid nitrogen and stored at −80 °C until extraction. Pigments were extracted from the frozen pellet using 100% acetone (Wako, Japan) with ultrasonic treatment for 10 min in an ice-water bath. The samples were then centrifuged at 15,000 rpm for 5 min at 4 °C. The supernatant was filtered through a 0.45-µm hydrophilic PTFE filter (SLPT0445NL, Hawach Scientific, IL, USA) and subjected to HPLC analysis. The HPLC method was based on the protocol of Nagao (Nagao et al. 2013b), which was a modified version of the method originally described by Zapata (Zapata M et al. 2000). Solution A contained 0.25 mol L⁻¹ pyridine, adjusted to pH 5.0 with acetic acid. The pigment composition was analyzed using a Shimadzu HPLC system equipped with LC-20AD pumps and an SPD-M20A photodiode array detector. Separation was performed using an Inertsil C8 reversed-phase column (5020-01228, GL Sciences, Japan). The injection volume for each sample was 20 µL. Standards for chlorophyll *a*, fucoxanthin, diadinoxanthin, and diatoxanthin were purchased from DHI (Denmark) and analyzed in parallel to construct calibration curves. Xanthophyll de-epoxidation state (DES) was calculated as the ratio of diatoxanthin to the sum of diadinoxanthin and diatoxanthin.

### Preparation of thylakoid membranes from Chaetoceros gracilis

*Chaetoceros gracilis* cells were disrupted by bead beating using 0.5 g of 0.1 mm glass beads. Disruption consisted of three 10 second pulses with a buffer 20 mM Tricine-KOH pH 7.6 0.4 M sorbitol 10% (w/v) PEG-6000 10 mM EDTA. The supernatant was collected by centrifugation at 21,500 × g for 5 min at 4 °C. The resulting pellet was resuspended in 1 mL of wash buffer (20 mM Tricine-KOH, pH 7.6; 0.4 M sorbitol; 5 mM MgCl₂; 2.5 mM EDTA) and subjected to low-speed centrifugation (50 × g 5 min 4 °C) to remove the cellular debris. The middle layer, containing purified thylakoid membranes, was carefully collected without disturbing the upper and lower impurities and centrifuged again at 21,500 × g for 5 min at 4 °C to pellet the thylakoid membranes. The thylakoid pellet was resuspended in solubilization buffer (50 mM imidazole-HCl, pH 7.0 at 4 °C; 20% glycerol).

### Combined CN/SDS-2D-PAGE and MS analyses

CN–PAGE was performed with modifications to published procedures (Järvi et al. 2011; Kameo et al. 2021). Thylakoid membranes were solubilized by solubilization buffer (50 mM imidazole-HCl, pH 7.0 at 4 °C; 20% glycerol) and adjusted to a final concentration of 1 μg Chl *a* + *c*. For DOC-based CN-PAGE, an equal volume of 2% (w/v) α-DM was added to solubilize the membranes. After incubation on ice for 30 minutes, the samples were centrifuged to remove the insoluble debris. The resulting supernatant with 5 μg of Chl (in 10 μL) was loaded per well in the CN-PAGE gel. Two types of the CN-PAGE gel consist of 3.5 % acrylamide (37.5:1) for the stacking gel and a 4%–13%/6%–14% acrylamide (37.5:1) gradient for the separation gel. Electrophoresis was performed at 4 °C using the myPowerII 300 system (AE-8135, ATTO, Japan) at 300 V and 2 mA for 180 minutes. Selected bands from CN-PAGE were excised from the gels and identified by liquid chromatography–tandem MS.

For subsequent SDS-PAGE analysis, the gel strips were incubated at room temperature in solubilization buffer (1% SDS and 50 mM DTT) for 30 minutes. Samples were then loaded onto a stacking gel (pH 6.8) and a separation gel composed of 14% acrylamide (37.5:1), 6 M urea, and pH 8.6. Electrophoresis was carried out using the Laemmli system. Gels were initially run at 250 V and 100 mA until the tracking dye reached the bottom of the marker well. Precision Dual Color Standards (#1610374, Bio-Rad, CA, USA) were then added, and electrophoresis was performed at 250 V for an additional 50 minutes (total ∼90 min). After electrophoresis, gels not used for immunoblotting were stained with Oriole fluorescent gel stain (#1610496, Bio-Rad, CA, USA) and imaged using a ChemiDoc Touch Imaging System (Bio-Rad).

### Immunoblotting analysis

Diatom cells were solubilized in SDS-PAGE sample buffer (62.5 mM Tris–HCl (pH 6.8), 2.5% (w/v) SDS, 10% (w/v) glycerol, 2.5% (v/v) 2-mercaptoethanol trace bromophenol blue). The solubilization was performed by heating at 37 °C for 30 minutes followed by a 30-minute incubation period at room temperature. The lysates were then centrifuged, and the protein concentration was adjusted to 30 μg per 20 μL. For the detection of Lhcx1 or D1 protein, SDS-PAGE was performed using a 15% acrylamide separation gel and a 4.5% stacking gel. 1.25/2.5/5 μg of the protein was loaded per lane. The anti–Lhcx1 peptide antibody was raised against the synthetic peptide “KEEYAPGDLRFDPFGLMP” for Lhcx1 protein. Antibody against the carboxy-terminal fragment of D1 (AS05084) was purchased from Agrisera. After electrophoresis, the gel was desalted and equilibrated in electron transfer buffer (25 mM Tris, 192 mM glycine, 20% (v/v) methanol, 0.05% (w/v) SDS), and proteins were transferred to an Immobilon-P PVDF membrane (Millipore, Billerica, MA, USA). The membrane was incubated with a 1:5000 dilution of the anti–Lhcx1 antibody. Immunoreactive proteins were visualized using the Amersham ECL Prime Western Blotting Detection Reagent (Cytiva, RPN2236) and imaged with the ChemiDoc Touch Imaging System (Bio-Rad). For the detection of Lhcx1 on the CN-PAGE, the CN-PAGE after electrophoresis was prepared with the same protocol as SDS-PAGE.

### RNA extraction and RT-qPCR analysis

Cells were collected by centrifugation and subjected to RNA extraction. Total RNA was extracted using the RNA Easy Fast Plant Tissue Kit (DP452, Tiangen) according to the manufacturer’s instructions. Total RNA was treated with gDNA Eraser using the PrimeScript RT reagent Kit (Perfect Real Time) (TaKaRa, RR037A) to remove genomic DNA. Reverse transcription was then performed using the same kit, following the manufacturer’s instructions. Each qPCR reaction consisted of 5 μL of SsoAdvanced Universal SYBR Green Supermix (Bio-Rad), 2 μL of nuclease-free water, 2 μL of cDNA template, and 0.5 μL each of forward and reverse primers (10 μM each before mixing), in a total volume of 10 μL. RT-qPCR was performed using the CFX96 Real-Time PCR Detection System (Bio-Rad). The β-tubulin and β-actin genes were used as reference genes. The ΔΔCt method was used for relative quantification of the transcript abundance of target genes across samples. Primer sequences for CgLhcx1 and the reference genes used in qRT-PCR analysis are reported in Supplementary Data 1.

### Phylogenetic analysis

LHC sequences were collected from genomic and transcriptomic data available in the PhycoCosm and EukProt databases following the previously described method (Kumazawa and Ifuku 2024), in addition to our earlier dataset (Kumazawa et al. 2022; Kumazawa and Ifuku 2024). The compiled dataset includes LHC sequences from centric diatoms, including *T*, *pseudonana*, *Thalassiosira oceanica*, and *C*, *Meneghiniana*, as well as from pennate diatoms such as *P. tricornutum*, *Fragilariopsis cylindrus*, *Fistulifera solaris*, and *Pseudo-nitzschia multistriata*. Redundant LHC protein sequences were removed using CD-HIT (Fu et al. 2012), and the remaining sequences were aligned using MAFFT-einsi v7.525, followed by manual curation (Katoh and Standley 2013). The curated dataset was realigned and trimmed using ClipKit v2.3.0 in the smart-gap mode to eliminate unaligned regions (Steenwyk et al. 2020). Phylogenetic trees were inferred using IQ-TREE v2.3.6 with the Q.pfam+I+R7 model, selected by ModelFinder from among the LG and Q.pfam-based models (Kalyaanamoorthy et al. 2017; Minh et al. 2020, 2021), and visualized using iTOL v6 (Letunic and Bork. 2024). Ultrafast bootstrap approximation (UFBoot) and SH-aLRT support tests were performed with 1000 replicates each (Guindon et al. 2010; Hoang et al. 2018), and aBayes support was also assessed (Anisimova et al. 2011). Subfamily annotations of LHCs in the phylogenetic trees were based on our previous studies (Kumazawa and Ifuku 2024) and were illustrated using Inkscape.

### Statistical analysis

All data in the text, tables, and figures are reported as mean ± standard deviation (SD), unless otherwise indicated. We used two-way ANOVA followed by Dunnett’s multiple comparisons test to determine the statistical significance of differences among genotypes. All statistical analyses were performed and visualized using GraphPad (https://www.graphpad.com/).

### Accession numbers

For *Chaetoceros gracilis* genes, Gene IDs from the v1 genome assembly available at our *Chaetoceros gracilis* genome database (https://chaetoceros.nibb.ac.jp/) are provided as follows: Lhcf2, g4294.t1; Lhcx1, g4948.t1; *β*-actin, g9224.t1; *β*-tubulin, g11207.t1.

## Acknowledgments

This work was supported in part by JSPS KAKENHI grant Nos. JP25KJ1660 (J.X.), JP23H02347 (K.I.), JP24H02081 (K.I.), and a grant from the Institute for Fermentation, Osaka, Japan (K.I.). We thank Dr. Shoko Tsuji of Kyoto University for her assistance with the pigment analysis and Dr. Noriko Ishikawa for her help with the plasmid construction.

## Author Contributions

K.I. conceived the project; J.X. carried out all physiological and biochemical analyses; J.X. and M.K. performed the phylogenetic analysis; J.X. drafted the original manuscript; J.X., M.K., and K.I. revised and finalized the manuscript; all authors participated in discussing the results.

