## Supplementary Fig. for "Lhcf2 in the peripheral antenna is essential for non-photochemical quenching and Lhcx1 accumulation in the diatom *Chaetoceros gracilis*"

*Corresponding Authors:

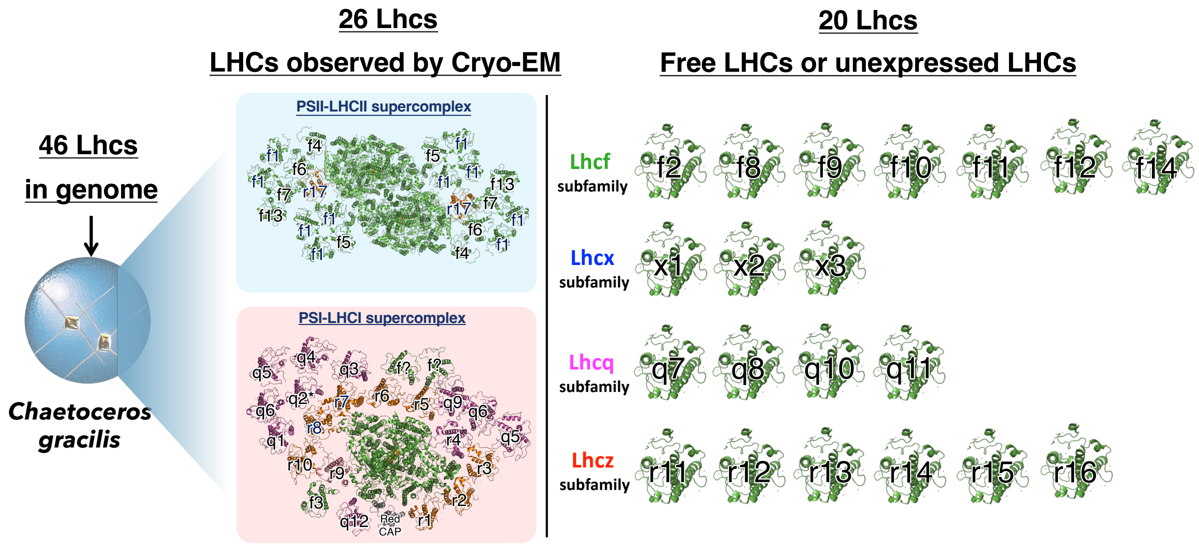


**Supplementary Fig. 1 | Unresolved LHC proteins in *C. gracilis*.**

Schematic representation of the PSI–LHCI and PSII–LHCII supercomplexes. The right panel details LHC proteins that were not resolved in the current cryo-EM structure, categorized four subfamilies: Lhcf, Lhcx, Lhcq, and Lhcz.

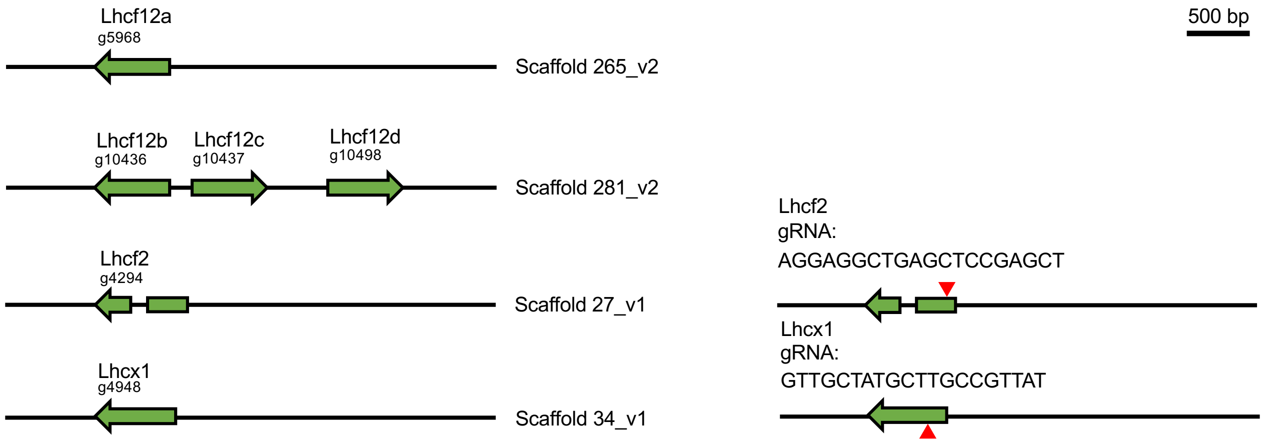


**Supplementary Fig. 2 | Gene schematics and gRNA design for CRISPR-/Cas9-mediated gene knockout of lhcf2 and lhcx1 genes.**

Genomic loci of the target genes, *lhcf12*, *lhcf2* and *lhcx1*, in *C. gracilis*. Coding sequences are represented by arrows, with corresponding gene names labeled above each gene structure. The specific gRNA sequences designed to target *hcf2* and *hcx1* are shown to the right of each gene diagram. Red arrowheads indicate target sites of gRNA utilized for the CRISPR/Cas9-mediated genome editing strategy.

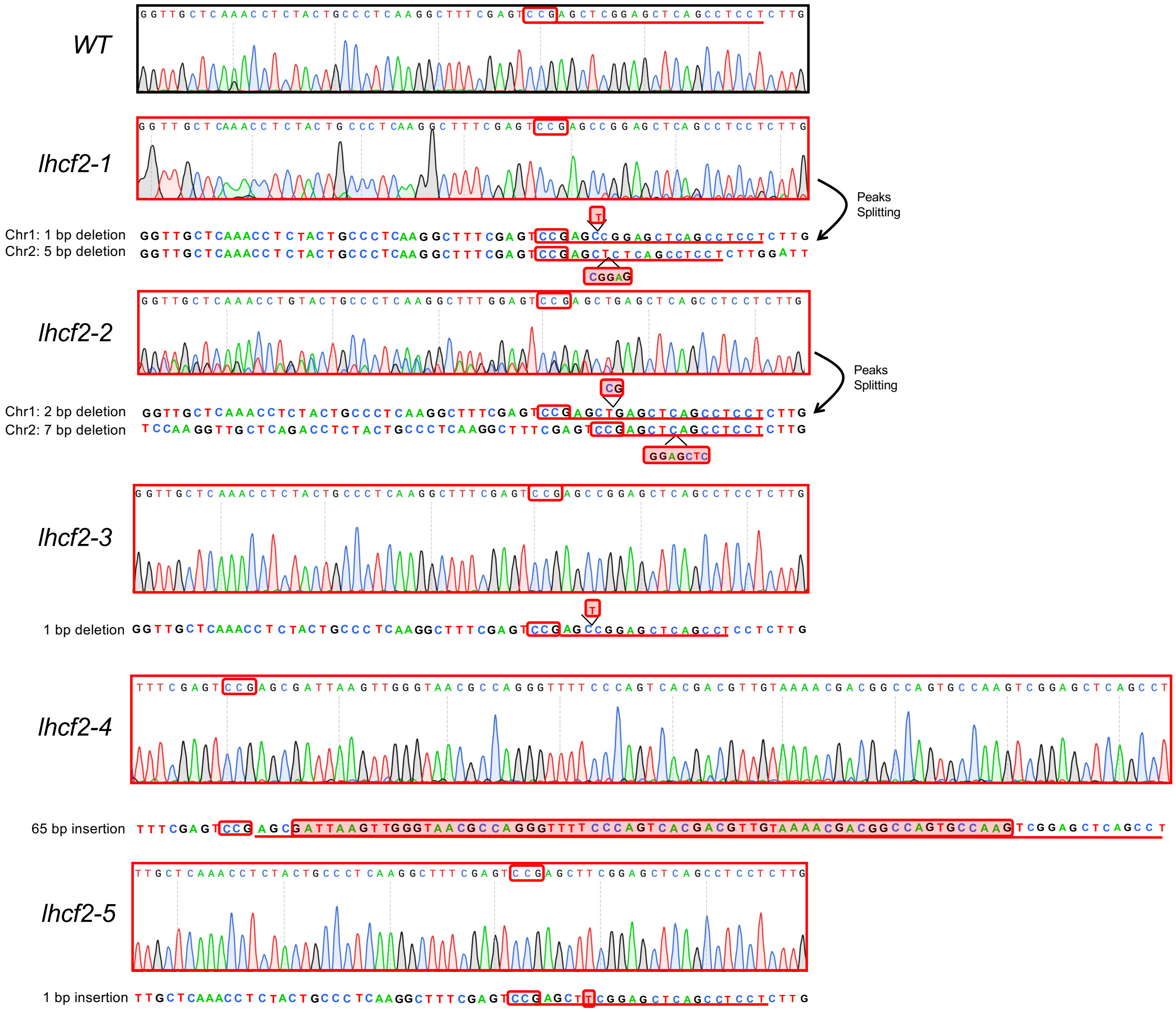


**Supplementary Fig. 3 | Genotypes of *lhcf2* mutants.**

Sanger sequencing chromatograms comparing WT and five independent *lhcf2* mutants are shown. CRISPR/Cas9-mediated genome editing resulted in insertions and deletions (indels) at the target site within the *lhcf2* locus. In the *lhcf2-1* mutant, a 1-bp deletion was detected on chromosome 1 and a 5-bp deletion on chromosome 2. In the *lhcf2-2* mutant, a 2-bp deletion on chromosome 1 and a 7-bp deletion on chromosome 2 were observed. In *lhcf2-3*, *lhcf2-4*, and *lhcf2-5* mutants, a 1-bp deletion, a 65-bp insertion, and a 1-bp insertion were detected, respectively. Red boxes indicate the positions of deleted or inserted bp, and peak splitting in the chromatograms indicates mixed alleles or frameshift-induced sequence divergence between the two alleles. These indels are predicted to cause frameshifts and premature stop codons, resulting in loss of Lhcf2 protein synthesis.

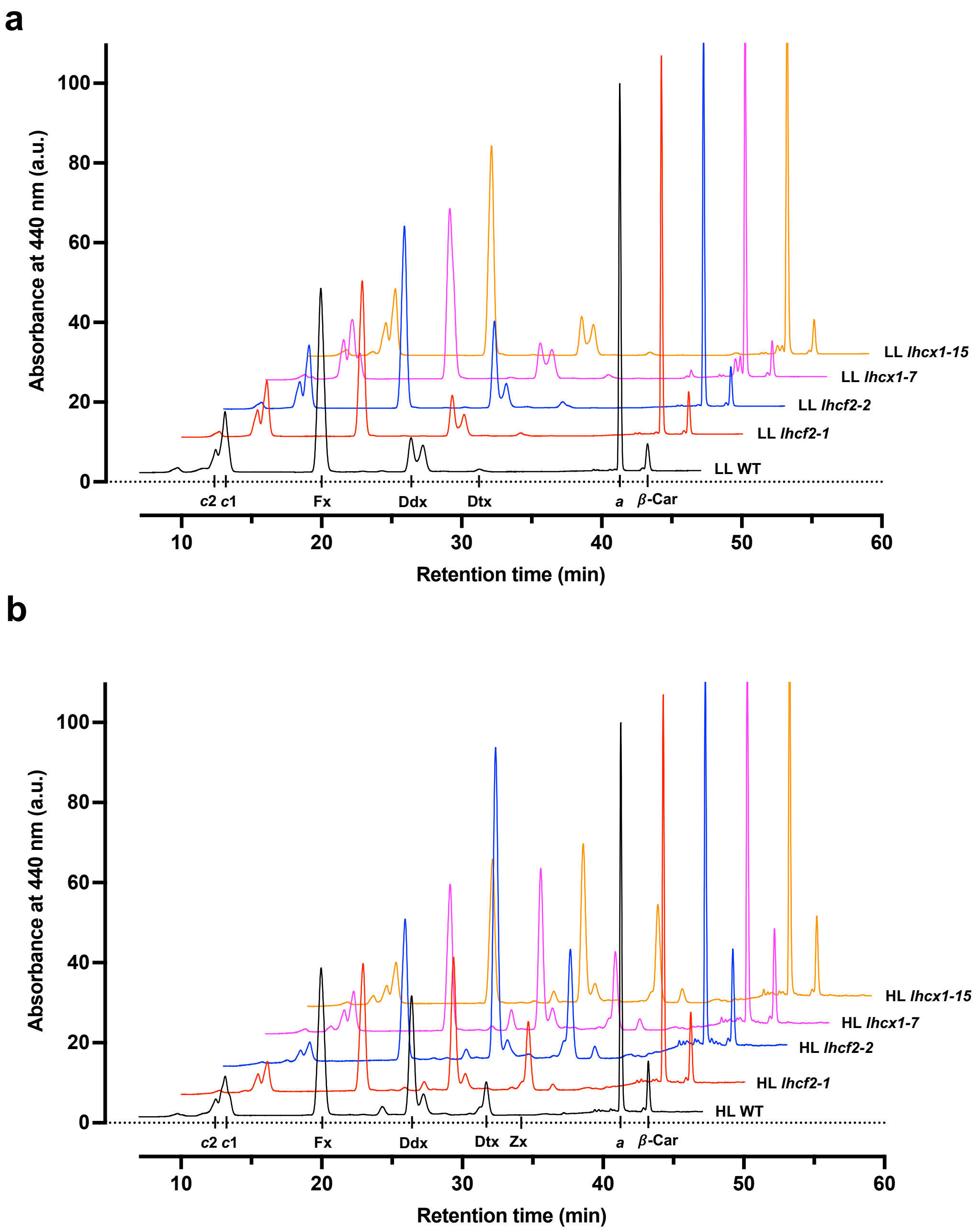


**Supplementary Fig. 4 | Pigment analysis using HPLC between WT and *lhcf2* mutants.**
HPLC traces for pigment extracts obtained from WT, *lhcx1*, and the *lhcf2* mutant under LL (a) and HL (b) conditions, analyzed by HPLC monitored at 440 nm and normalized to the Chl *a* peak. Peaks correspond to Chl *c*, Fx, Ddx, Dtx, Zx, Chl *a*, and β-Car. Data in each figure represent mean with biological replicates of n = 3.

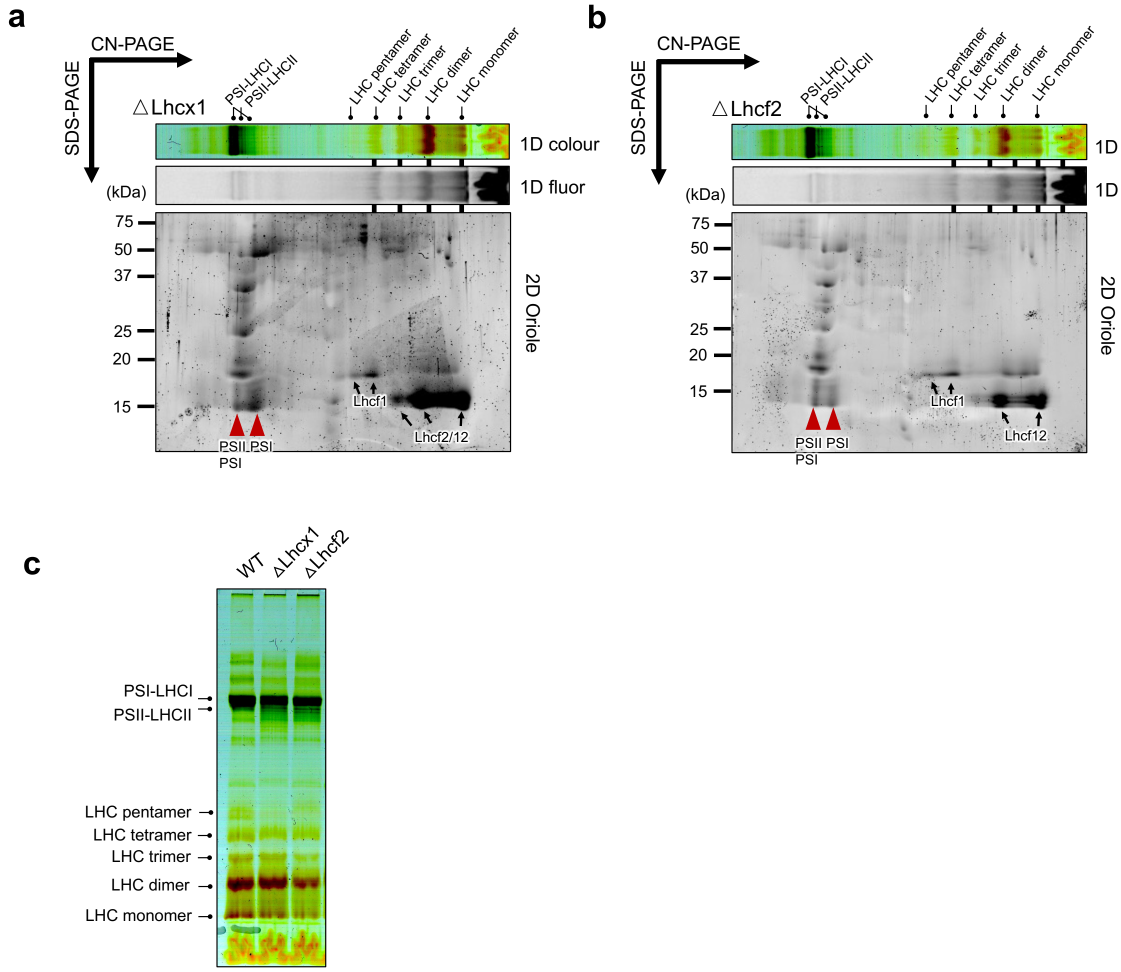


**Supplementary Fig. 5 | Comparative CN/2D-PAGE WT and NPQ mutants.**

**a, b**) Thylakoid membranes isolated from WT (a) and *hcf2* (b) mutant strains were solubilized using $\alpha$-DDM and sodium deoxycholate (DOC) separated by CN-PAGE (6–14% gradient) in the first dimension, and by SDS-PAGE in the second dimension. Fluorescence, pigment, and Oriole-stained profiles are above each corresponding 2D gel. Red arrowheads indicate the positions of PSI–LHCI or PSII–LHCII supercomplexes. **c**) CN-PAGE (6–14% gradient) analysis of WT, *lhcx1,* and *lhcf2* mutants.

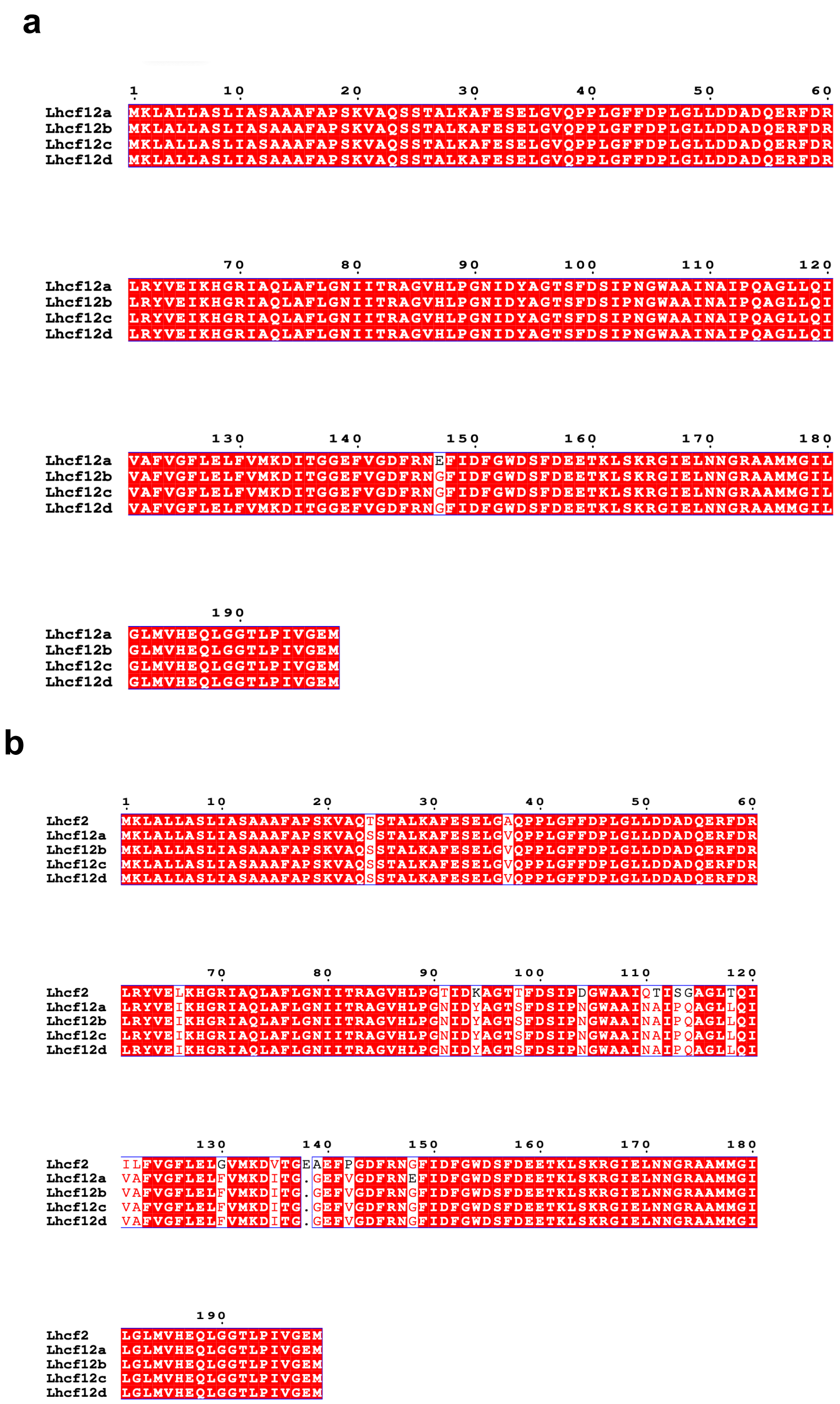


**Supplementary Fig. 6 | Amino acid alignments of LHCs analyzed in this study.**
**a)** multiple sequence alignment of Lhcf12a, Lhcf12b, Lhcf12c, and Lhcf12d reveals nearly identical sequences, with only a single amino acid substitution distinguishing Lhcf12a from the other three isoforms. **b)** multiple sequence alignment of Lhcf2 with the four Lhcf12 variants shows extensive sequence similarity, particularly in the C-terminal region, which complicates their discrimination in MS-based proteomic analysis. Conserved residues are highlighted in red, and mismatches are marked in white with black borders.

**a**

| Number | m/z (mi) | m/z (av) | Modifications | Start | End | Missed Cleavages | Sequence |
|---|---|---|---|---|---|---|---|
| 1 | 736.4240 | 736.8916 | | 33 | 37 | 0 | (R)Abu-YVELK(H) |
| 1 | 791.4522 | 791.9338 | | 28 | 32 | 1 | (R)Abu-FDRLR(Y) |
| 1 | 957.5112 | 958.0683 | | 137 | 144 | 0 | (R)Abu-GIELNNGR(A) |
| 1 | 1005.6091 | 1006.2407 | | 31 | 37 | 1 | (R)Abu-LRYVELK(H) |
| 1 | 1086.6055 | 1087.2745 | | 33 | 40 | 1 | (R)Abu-YVELKHGR(I) |
| 1 | 1113.6123 | 1114.2570 | | 136 | 144 | 1 | (K)Abu-RGIELNNGR(A) |
| 1 | 1192.6684 | 1193.3968 | | 54 | 64 | 0 | (R)Abu-AGVHLPGTIDK(A) |
| 1 | 1514.9053 | 1515.8505 | | 41 | 53 | 0 | (R)Abu-IAQLAFLGNIITR(A) |
| 1 | 1524.6965 | 1525.6245 | | 104 | 116 | 0 | (K)Abu-DVTGEAEFPGDFR(N) |
| 1 | 1865.0868 | 1866.2334 | | 38 | 53 | 1 | (K)Abu-HGRIAQLAFLGNIITR(A) |
| 1 | 1991.8658 | 1993.1063 | | 117 | 132 | 0 | (R)Abu-NGFIDFGWDSFDEETK(L) |
| 1 | 2320.0768 | 2321.5206 | | 117 | 135 | 1 | (R)Abu-NGFIDFGWDSFDEETKLSK(R) |
| 1 | 2603.5032 | 2605.1182 | | 41 | 64 | 1 | (R)Abu-IAQLAFLGNIITRAGVHLPGTIDK(A) |
| 1 | 2653.3584 | 2655.2990 | | 145 | 169 | 0 | (R)Abu-AAMMGILGLMVHEQLGGTLPIVGEM(-) |
| 1 | 2669.3533 | 2671.2983 | 1Oxidation | 145 | 169 | 0 | (R)Abu-AAMMGILGLMVHEQLGGTLPIVGEM(-) |
| 1 | 2685.3482 | 2687.2977 | 2Oxidation | 145 | 169 | 0 | (R)Abu-AAMMGILGLMVHEQLGGTLPIVGEM(-) |
| 1 | 2701.3432 | 2703.2971 | 3Oxidation | 145 | 169 | 0 | (R)Abu-AAMMGILGLMVHEQLGGTLPIVGEM(-) |
| 1 | 2717.3381 | 2719.2965 | 4Oxidation | 145 | 169 | 0 | (R)Abu-AAMMGILGLMVHEQLGGTLPIVGEM(-) |
| 1 | 3018.4367 | 3020.2995 | 1Abu->Acetyl | 1 | 27 | 0 | (-)Abu-FESELGAQPPLGFFDPLGLLDDADQER(F) |
| 1 | 3061.4789 | 3063.3683 | | 1 | 27 | 0 | (-)Abu-FESELGAQPPLGFFDPLGLLDDADQER(F) |
| 1 | 3412.4917 | 3414.6017 | | 104 | 132 | 1 | (K)Abu-DVTGEAEFPGDFRNGFIDFGWDSFDEETK(L) |
| 1 | 3436.6332 | 3438.7551 | 1Abu->Acetyl | 1 | 30 | 1 | (-)Abu-FESELGAQPPLGFFDPLGLLDDADQERFDR(L) |
| 1 | 3479.6754 | 3481.8239 | | 1 | 30 | 1 | (-)Abu-FESELGAQPPLGFFDPLGLLDDADQERFDR(L) |
| 1 | 3506.7990 | 3509.2382 | | 137 | 169 | 1 | (R)Abu-GIELNNGRAAMMGILGLMVHEQLGGTLPIVGEM(-) |
| 1 | 3522.7939 | 3525.2376 | 1Oxidation | 137 | 169 | 1 | (R)Abu-GIELNNGRAAMMGILGLMVHEQLGGTLPIVGEM(-) |
| 1 | 3538.7889 | 3541.2369 | 2Oxidation | 137 | 169 | 1 | (R)Abu-GIELNNGRAAMMGILGLMVHEQLGGTLPIVGEM(-) |
| 1 | 3554.7838 | 3557.2363 | 3Oxidation | 137 | 169 | 1 | (R)Abu-GIELNNGRAAMMGILGLMVHEQLGGTLPIVGEM(-) |
| 1 | 3570.7787 | 3573.2357 | 4Oxidation | 137 | 169 | 1 | (R)Abu-GIELNNGRAAMMGILGLMVHEQLGGTLPIVGEM(-) |

**b**

| Number | m/z (mi) | m/z (av) | Modifications | Start | End | Missed Cleavages | Sequence |
|---|---|---|---|---|---|---|---|
| 1 | 801.4213 | 801.8827 | | 148 | 153 | 0 | (K)Abu-ELQNGR(L) |
| 1 | 903.4894 | 904.0167 | | 25 | 32 | 0 | (K)Abu-ATDATLAR(Y) |
| 1 | 967.4956 | 968.0635 | | 35 | 42 | 0 | (R)Abu-EAELAHGR(V) |
| 1 | 1112.5946 | 1113.2634 | | 172 | 181 | 0 | (K)Abu-GIIEHLTSSA(-) |
| 1 | 1134.5426 | 1135.2260 | | 119 | 127 | 0 | (K)Abu-EEYAPGDLR(F) |
| 1 | 1222.6539 | 1223.3826 | | 25 | 34 | 1 | (K)Abu-ATDATLARYR(E) |
| 1 | 1286.6600 | 1287.4293 | | 33 | 42 | 1 | (R)Abu-YREAELAHGR(V) |
| 1 | 1531.8916 | 1532.9409 | | 43 | 56 | 0 | (R)Abu-VAMLAVIGFLVGEK(V) |
| 1 | 1547.8866 | 1548.9403 | 1Oxidation | 43 | 56 | 0 | (R)Abu-VAMLAVIGFLVGEK(V) |
| 1 | 1860.9888 | 1862.2119 | | 154 | 171 | 0 | (R)Abu-LAMLAAAGFLAQEAVDGK(G) |
| 1 | 1876.9837 | 1878.2113 | 1Oxidation | 154 | 171 | 0 | (R)Abu-LAMLAAAGFLAQEAVDGK(G) |
| 1 | 2223.0750 | 2224.5127 | | 99 | 118 | 0 | (R)Abu-AQEGWVDPADCPVDQPGLLK(E) |
| 1 | 2395.3166 | 2396.8753 | | 35 | 56 | 1 | (R)Abu-EAELAHGRVAMLAVIGFLVGEK(V) |
| 1 | 2411.3115 | 2412.8746 | 1Oxidation | 35 | 56 | 1 | (R)Abu-EAELAHGRVAMLAVIGFLVGEK(V) |
| 1 | 2460.3385 | 2461.8788 | 1Abu->Acetyl | 1 | 24 | 0 | (-)Abu-ASLEGIPGALPPVGIFDPLNLAEK(A) |
| 1 | 2471.1145 | 2472.8123 | | 128 | 147 | 0 | (R)Abu-FDPFGLMPEDPEEFDIMQTK(E) |
| 1 | 2487.1094 | 2488.8117 | 1Oxidation | 128 | 147 | 0 | (R)Abu-FDPFGLMPEDPEEFDIMQTK(E) |
| 1 | 2503.1044 | 2504.8111 | 2Oxidation | 128 | 147 | 0 | (R)Abu-FDPFGLMPEDPEEFDIMQTK(E) |
| 1 | 2503.3807 | 2504.9476 | | 1 | 24 | 0 | (-)Abu-ASLEGIPGALPPVGIFDPLNLAEK(A) |
| 1 | 2558.3395 | 2559.9655 | | 148 | 171 | 1 | (K)Abu-ELQNGRLAMLAAAGFLAQEAVDGK(G) |
| 1 | 2574.3344 | 2575.9649 | 1Oxidation | 148 | 171 | 1 | (K)Abu-ELQNGRLAMLAAAGFLAQEAVDGK(G) |
| 1 | 2869.5128 | 2871.3462 | | 154 | 181 | 1 | (R)Abu-LAMLAAAGFLAQEAVDGKGIIEHLTSSA(-) |
| 1 | 2885.5077 | 2887.3456 | 1Oxidation | 154 | 181 | 1 | (R)Abu-LAMLAAAGFLAQEAVDGKGIIEHLTSSA(-) |
| 1 | 3168.4653 | 3170.5659 | | 128 | 153 | 1 | (R)Abu-FDPFGLMPEDPEEFDIMQTKELQNGR(L) |
| 1 | 3184.4602 | 3186.5653 | 1Oxidation | 128 | 153 | 1 | (R)Abu-FDPFGLMPEDPEEFDIMQTKELQNGR(L) |
| 1 | 3200.4551 | 3202.5646 | 2Oxidation | 128 | 153 | 1 | (R)Abu-FDPFGLMPEDPEEFDIMQTKELQNGR(L) |
| 1 | 3253.5470 | 3255.6096 | | 99 | 127 | 1 | (R)Abu-AQEGWVDPADCPVDQPGLLKEEYAPGDLR(F) |
| 1 | 3259.7573 | 3261.7664 | 1Abu->Acetyl | 1 | 32 | 1 | (-)Abu-ASLEGIPGALPPVGIFDPLNLAEKATDATLAR(Y) |
| 1 | 3302.7995 | 3304.8352 | | 1 | 32 | 1 | (-)Abu-ASLEGIPGALPPVGIFDPLNLAEKATDATLAR(Y) |
| 1 | 3501.5865 | 3503.9092 | | 119 | 147 | 1 | (K)Abu-EEYAPGDLRFDPFGLMPEDPEEFDIMQTK(E) |
| 1 | 3517.5814 | 3519.9086 | 1Oxidation | 119 | 147 | 1 | (K)Abu-EEYAPGDLRFDPFGLMPEDPEEFDIMQTK(E) |
| 1 | 3533.5763 | 3535.9080 | 2Oxidation | 119 | 147 | 1 | (K)Abu-EEYAPGDLRFDPFGLMPEDPEEFDIMQTK(E) |

**Supplementary Fig. 7 | *In silico* prediction of tryptic peptides.**

Predicted tryptic peptides for Lhcf2 (a) and Lhcx1 (b) were generated under digestion settings: maximum of one missed cleavage, peptide mass range of 500–4000 Da, and minimum peptide length of five amino acids. A total of 18 peptides were computationally identified for use in proteomic analysis.

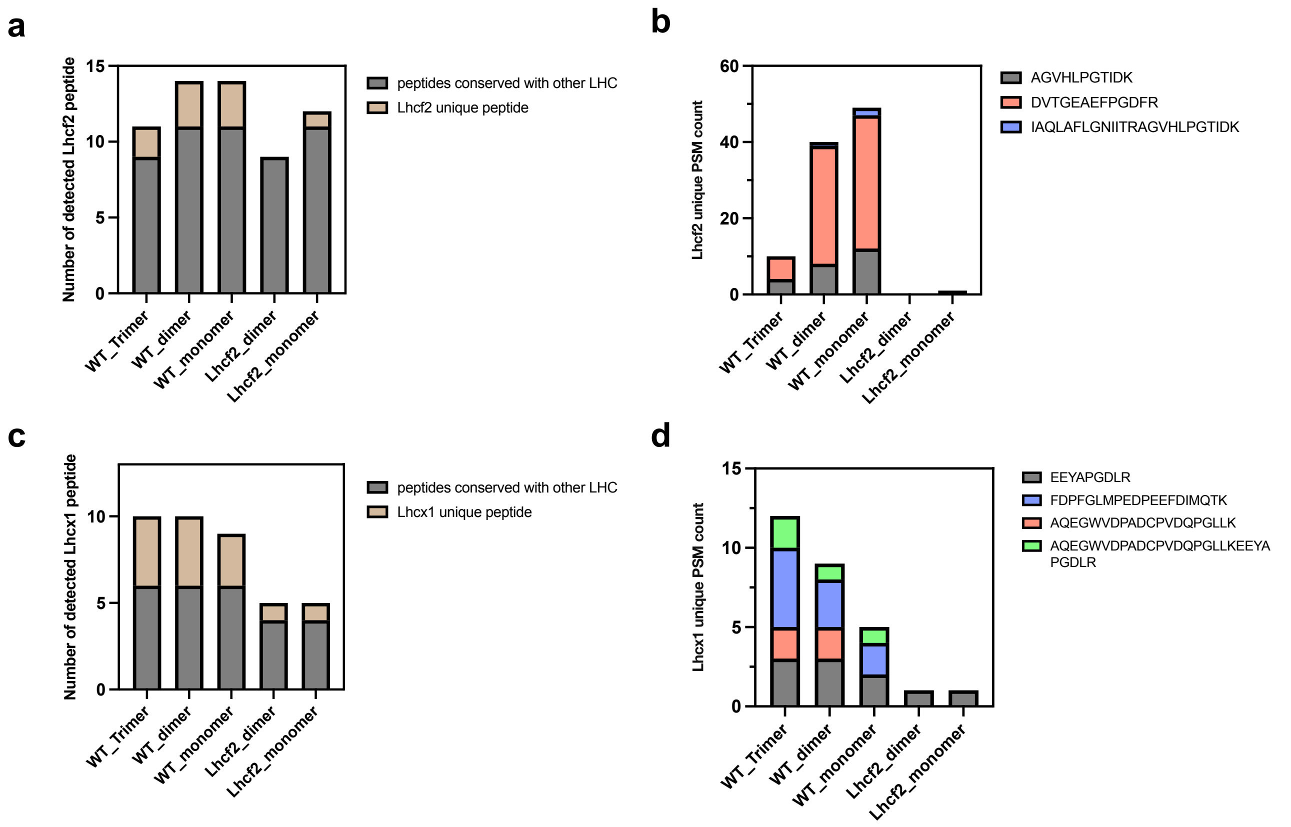


**Supplementary Fig. 8 | MS identification of CN-PAGE separated complexes.**

**a)** Number of Lhcf2 peptides identified across different LHC bands isolated from WT and *lhcf2* mutants. Unique peptides to Lhcf2 (brown) and peptides conserved with other LHCs (gray) are indicated. **b)** peptide spectrum match (PSM) counts of four Lhcf2-unique peptides detected in each CN-PAGE band. **c)** number of Lhcx1 peptides identified in different LHC bands from WT and *lhcf2* mutants. Unique peptides to Lhcx1 (brown) and peptides conserved with other LHCs (gray) are indicated. **d)** peptide spectrum match (PSM) counts of four Lhcx1-unique peptides detected in each CN-PAGE band.

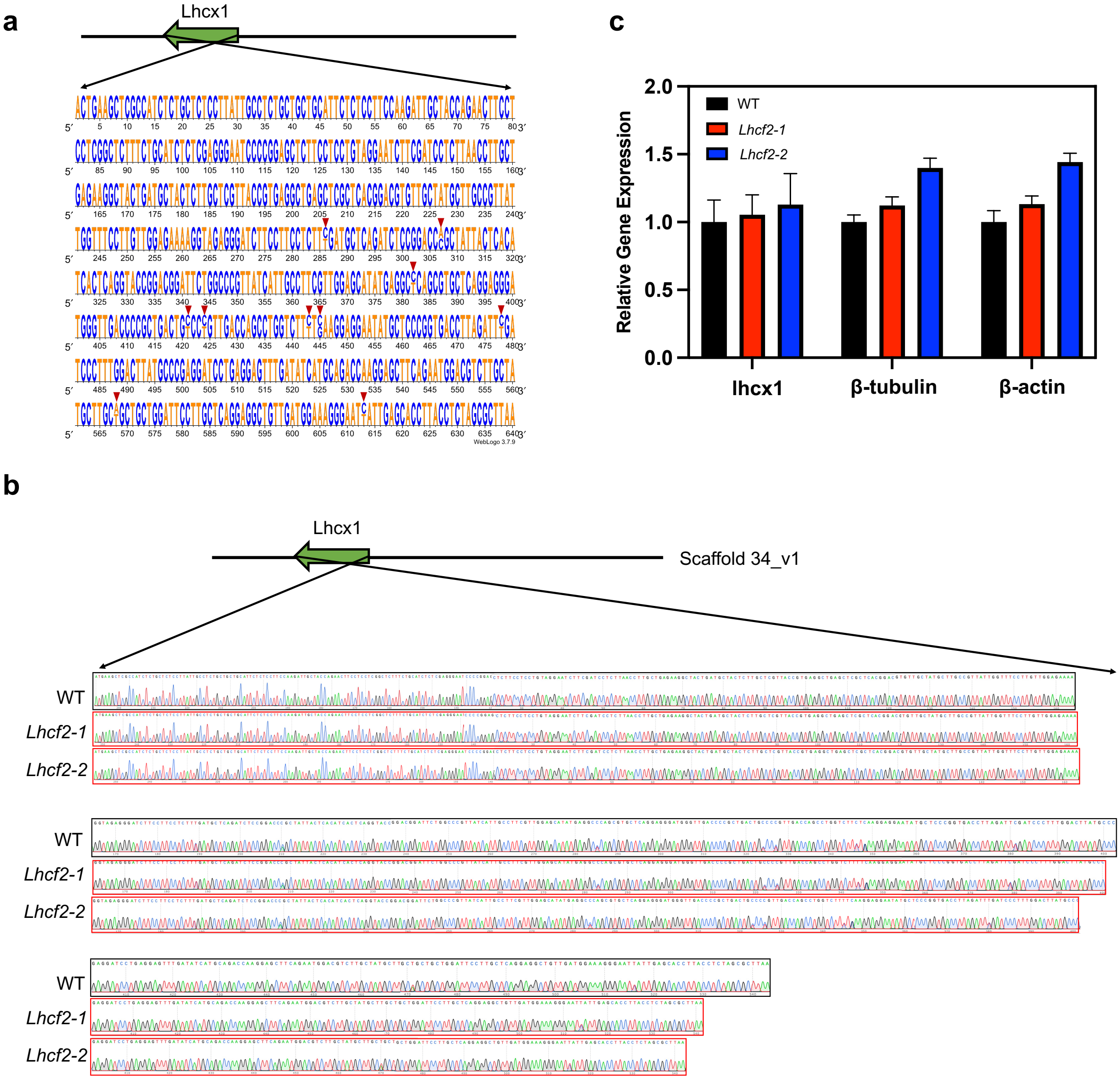


**Supplementary Fig. 9 | Confirmation of *Lhcx1* gene integrity and transcription in *lhcf2* mutants. a, b)** Schematic representation of the *lhcx1* genomic region (a) and Sanger sequencing chromatograms (b) of *lhcx1* PCR products from WT and *lhcf2* mutants. Double nucleotide peaks observed at certain positions (arrowed by red triangle. Although double peaks were observed at certain nucleotide positions, their similar intensity across all strains indicates natural heterozygosity rather than CRISPR-induced mutation. No mutation was detected in the *Lhcx1* coding sequence of the *lhcf2* mutants, confirming its intact genetic structure. **c)** Quantitative RT-qPCR analysis of lhcx1 transcript levels, normalized *to β-tubulin* and *β-actin* in WT and *lhcf2* mutants. Data represent mean ± SEM with biological replicates of n = 3-4.

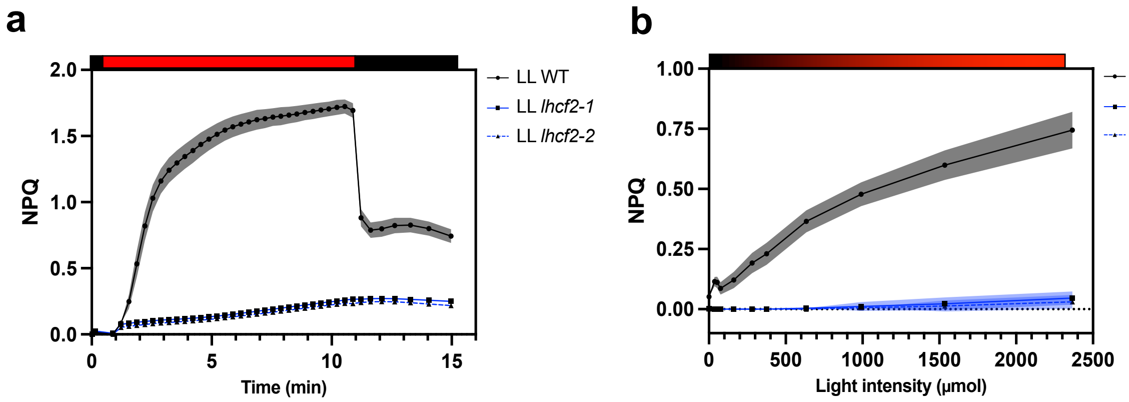


**Supplementary Fig. 10 | NPQ characteristics measured under red actinic light.**
Induction kinetics (a) and NPQ light response curve (b) for WT and *lhcf2* mutants grown under LL conditions.

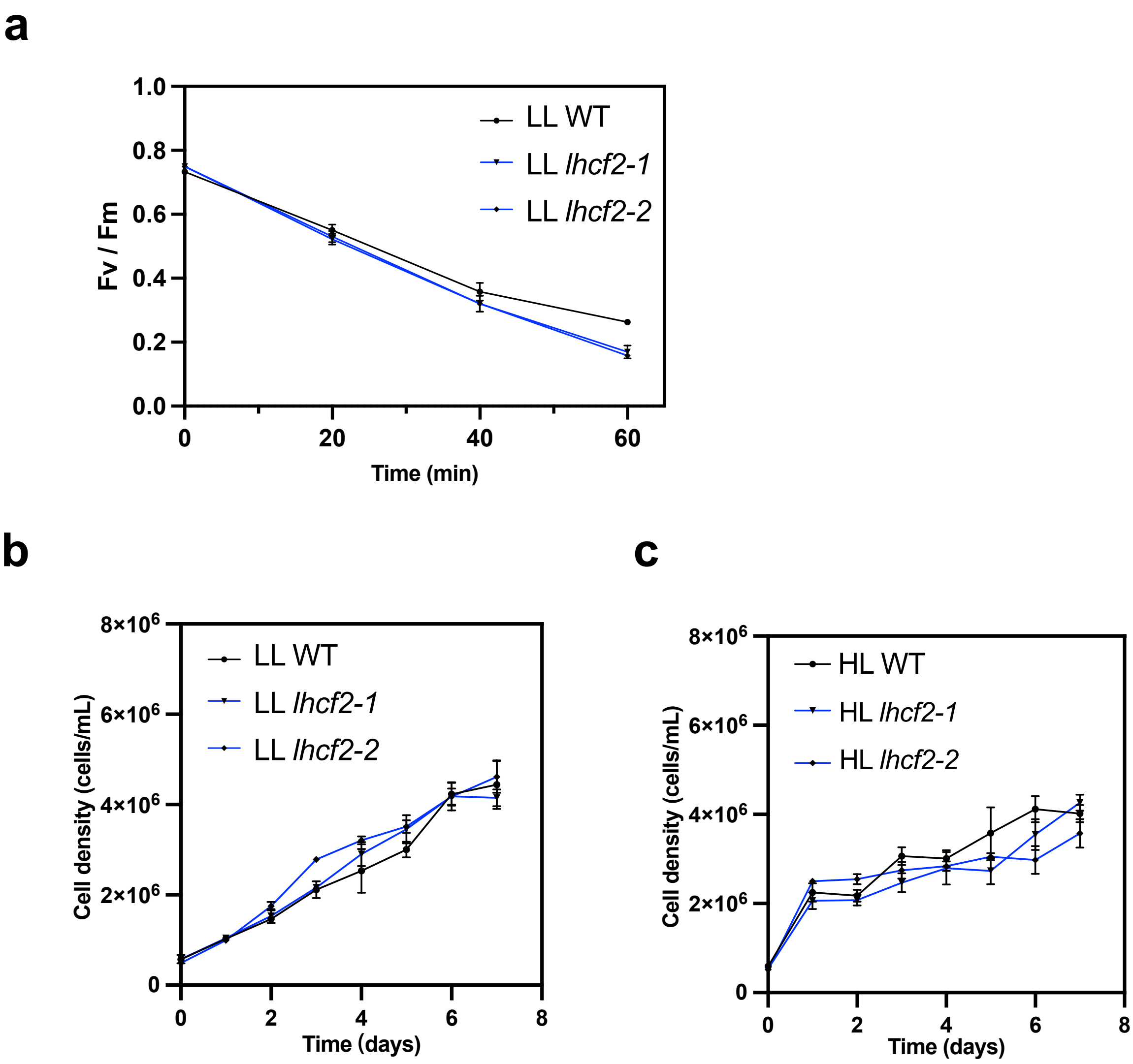


**Supplementary Fig. 11 | Growth rates and photoinhibition susceptibility analysis in WT and *lhcf2* mutants.**

**a**, **b**) Growth curve analysis between WT and *lhcf2* mutants grown under LL (a) and HL (b) conditions. **c)** Photoinhibition measurement of WT and *lhcf2* mutants. Data in each figure represent mean ± SD *n* = 3–4 biological replicates.

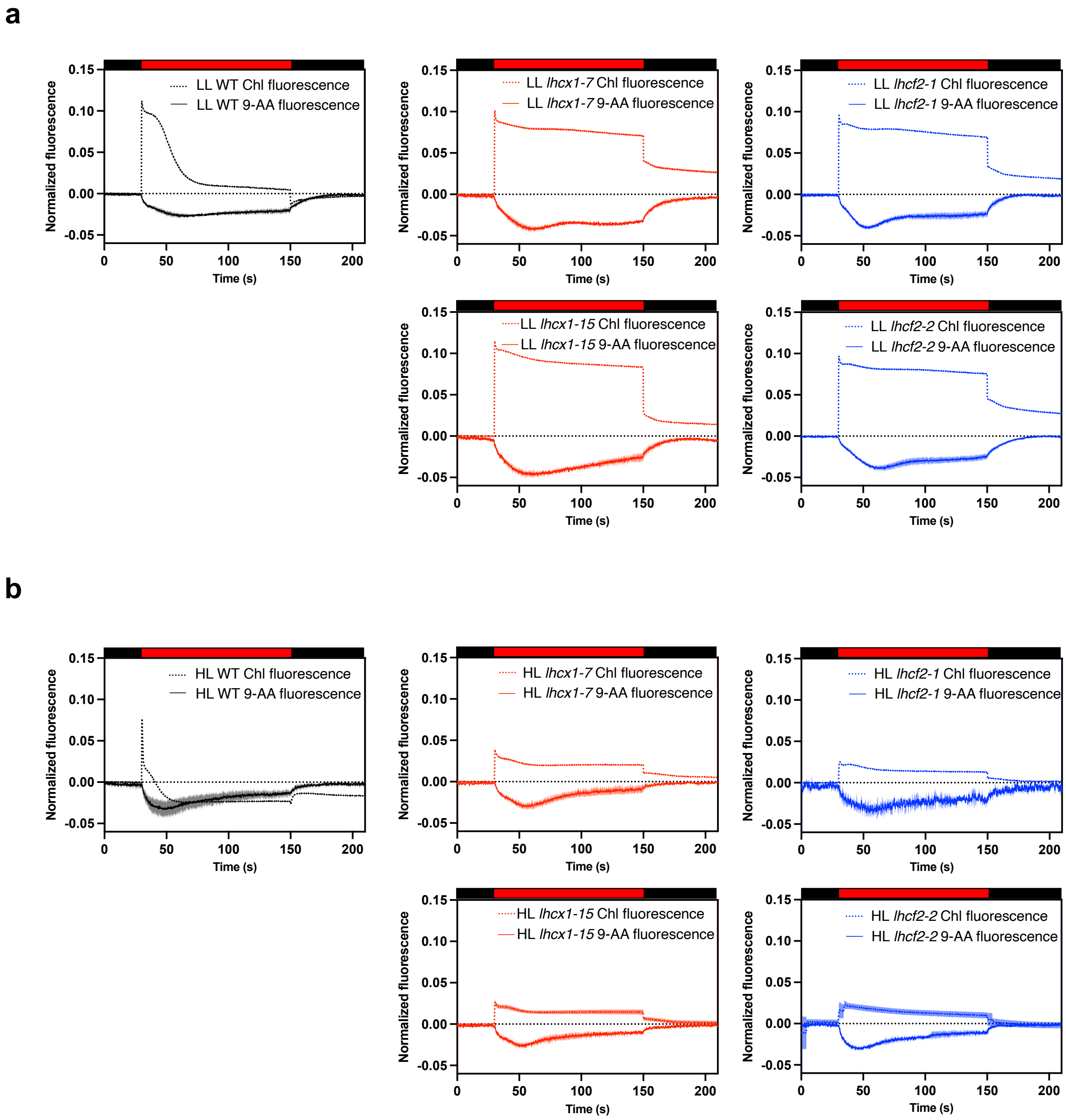


**Supplementary Fig. 12 | Characterization of the trans-thylakoid pH gradient (ΔpH) in *C. gracilis* living cells.**

Chlorophyll fluorescence transient and 9-aminoacridine (9-AA) fluorescence quenching kinetics WT, *lhcx1*, and *lhcf2* mutants grown under LL (a) and HL (b) conditions. Chlorophyll fluorescence and 9-AA quenching, which reports on ΔpH formation, were measured using a DUAL-PAM 100 fluorometer. Red box actinic light (1,400 μmol photons $m^{-2}$ $s^{-1}$) and the black box. Representative traces are shown from 2–3 biological replicates with similar results.

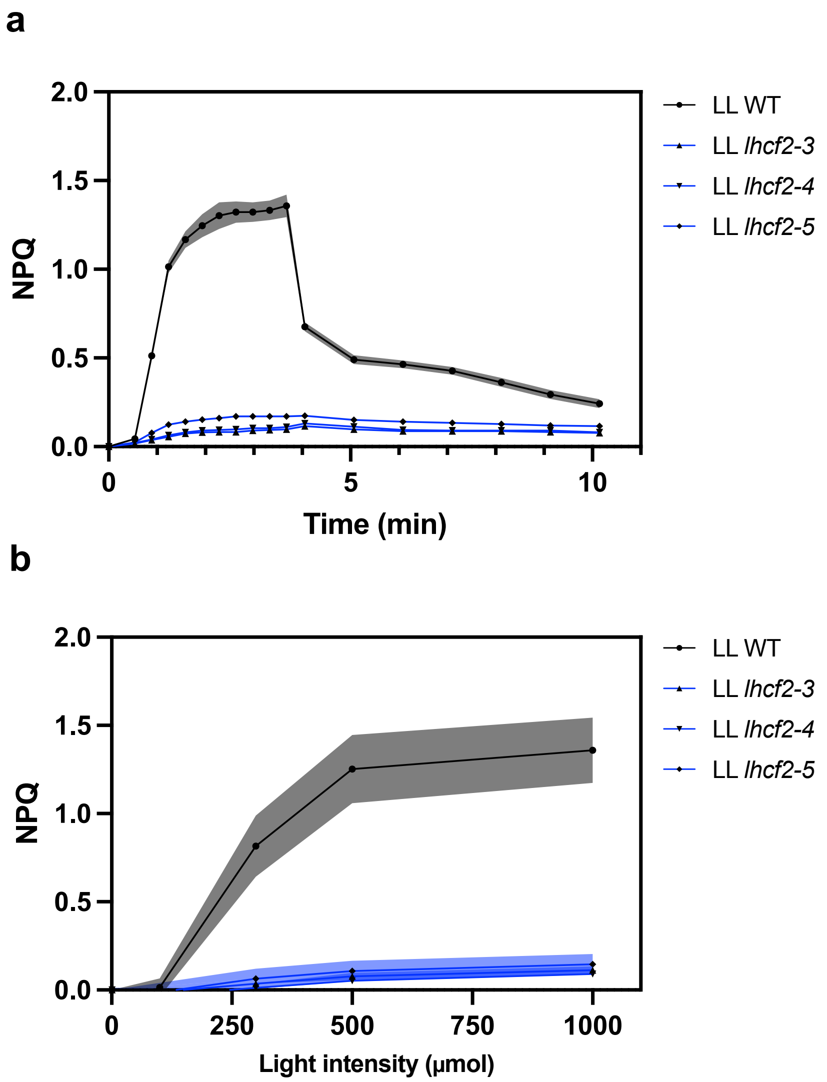


**Supplementary Fig. 13 | Confirmation of NPQ deficiency in independent *lhcf2-3,4,5* mutants.** Induction kinetics curve (a) and NPQ light response curve (b) measured by blue actinic light among WT and *lhcf2* mutants grown under LL conditions.

**Supplementary Data 1.**
Sequences of oligonucleotide primers used in this study.

**CRISPR sgRNA of *Lhcf2***

| | |
|---|---|
| Lhcf2_98-120_RGR | AGGAGGCTGAGCTCCGAGCT |

**Mutant genotyping PCR and sequencing primer**

| | |
|---|---|
| Primer of *Lhcf2*_Fw (sequencing primer) | TACTTACCAGCCATCGGGGA |
| Primer of *Lhcf2*_Rv | GGTATCCCATACGCTGTGGT |
| Primer of *Lhcx1*_F1_Fw | AGCTGTCCAACCTGCTTGAT |
| Primer of *Lhcx1*_ F1_Rv | TACTCCACACAACAGCAGCA |
| Primer of *Lhcx1*_F2_Rv (sequencing primer) | GAAGGAAGATCCCTCTACCT |
| Primer of *Lhcx1*_F3_Rv (sequencing primer) | AGAACTTCCTCCTCGGCTCT |

**RT-qPCR**

| | |
|---|---|
| RT-qPCR primer of *Lhcx1*_Fw | GCCATCTCTGCTCTCCTTATT |
| RT-qPCR primer of *Lhcx1*_Rv | AAGAGGATCGAAGATTCCTACA |
| RT-qPCR primer of *β-tubulin*_Fw | GGAGCTGAGCTTGTTGATTC |
| RT-qPCR primer of *β-tubulin*_Rv | ACCAGCTCCAGTTCCTCCTC |
| RT-qPCR primer of *β-actin*_Fw | CATACATGGCGGGGACAT |
| RT-qPCR primer of *β-actin*_Rv | GCTCACCCCGTCCTTCTTA |

*Sequences are shown from 5' to 3'.
